# Charge Interactions in a Highly Charge-depleted Protein

**DOI:** 10.1101/2020.06.26.173278

**Authors:** Stefan Hervø-Hansen, Casper Højgaard, Kristoffer Enøe Johansson, Yong Wang, Khadija Wahni, David Young, Joris Messens, Kaare Teilum, Kresten Lindorff-Larsen, Jakob Rahr Winther

**Author notes:** These authors contributed equally to this work (Stefan Hervø-Hansen, Casper Højgaard).

## Abstract

Interactions between charged residues are difficult to study because of the complex network of interactions found in most proteins. We have designed a purposely simple system to investigate this problem by systematically introducing individual and pairs of charged and titratable residues in a protein otherwise free of such residues. We used constant pH molecular dynamics simulations, NMR spectroscopy, and thermodynamic double mutant cycles to probe the structure and energetics of the interaction between the charged residues. We found that the partial burial of surface charges contributes to a shift in p*K*_a_ value, causing an aspartate to titrate in the neutral pH range. Additionally, the interaction between pairs of residues was found to be highly context dependent, with some pairs having no apparent preferential interaction, while other pairs would engage in coupled titration forming a highly stabilized salt bridge. We find good agreement between experiments and simulations, and use the simulations to rationalize our observations and to provide a detailed mechanistic understanding of the electrostatic interactions.

**Significance:** Electrostatic forces are important for protein folding and are favored targets of protein engineering. However, despite the many advances in the field of protein electrostatics, the prediction of changes in protein structure and function upon introduction or removal of titratable residues is still complicated. In order to provide a basic understanding of protein electrostatics we here characterize a highly charge-depleted protein and its titratable variants by a combination of NMR spectroscopy and constant pH molecular dynamics simulations. Our investigations reveal how strongly interacting residues engaged in salt bridging, can be characterized. Furthermore, our study may also enrich and facilitate the understanding of dehydration of salt-bridges and its potential effect on protein stability.

## Introduction

The most fundamental biochemical reactions in proteins like enzymatic catalysis [1], redox reactions [2], H^+^-transfer [3], and ion homeostasis [4], are governed by electrostatics. Electrostatic properties are substantially modulated by the protonation status of the titratable residues yielding positive and negative charges. By changing the solvent pH, we effectively change the electrostatic interactions which results in adaptation of protein function and/or protein structure. Consequently, from a protein engineering point of view, it is desirable to understand and predict the effects of changing the pH of the solution. Furthermore, the insertion/removal of titratable residues from proteins affects the p*K*_a_ values of other residues nearby which may impact on features like catalysis and structural stability. Over the last decades, a number of studies have addressed these problems utilizing theoretical, computational, and structure-based methods for the calculation of p*K*_a_ values and benchmarking against experimental studies. In a similar fashion, many studies have been conducted on various model systems with either artificial or biologically relevant salt bridges. The possibility of predicting the effect of a charge perturbation on stability and p*K*_a_ has resulted in many “rational modifications” by e.g. insertion of titratable residues causing charge-charge interactions [5], the filling of cavities by hydrophobic residues causing dehydration [6], and the insertion of polar residues causing hydrogen bonding [7]. Despite the many important findings by these studies, many supposedly rational substitutions cause unpredictable and complicated results. These have been attributed to either long-range electrostatic effects, the alteration of several interactions upon mutagenesis or denatured-state ensemble stabilization [8].

Due to the complication of predicting the outcome of inserting or removing titratable residues in proteins, we have turned to a model protein, that we have previously redesigned to be devoid of the ionizable amino acid residue types Asp, Glu, Arg, Lys, His, Tyr and (free) Cys [9]. This protein is based on the cellulose binding domain EXG:CBM [10], and is well folded, functional and stable, and makes it possible to reduce the complexity of the multiple electrostatic interactions seen in almost all model proteins studied hitherto. The charge-free protein consequently permitted us to study interactions between pairs of charged residues individually and at the same time measure the effect of partial burial of the charge pairs. We modeled the effect of introduced charges in the EXG:CBM system using discrete constant pH molecular dynamics (CpHMD) to provide pH-dependent atomistic structural details about the system and protonation states. To complement and validate our simulations experimentally, we followed the protonation state using nuclear magnetic resonance (NMR) spectroscopy, which, due to its high resolution, allows monitoring of site-specific effects. We further used double mutant cycles to estimate pH dependent coupling energies between charge pairs and to validate the protonation cycles obtained from simulations.

## Results & Discussion

### Structure Determination of a Charge-free Protein

We have previously described a variant of the cellulose binding domain protein EXG:CBM, from a *Cellulomonas fimi* cellulase, in which we substituted four ionizable amino acid residues with non-titratable residue types (K28M, D36Q, R68M and H90W) [9]. In most respects, this variant performed like the wild-type protein, however, protein yields from expression in *E. coli* were relatively low. For the current study, we identified a new charge-free variant, EXG:CBM^QQQW^, from a screen for higher protein yields. The new variant differs from the former by two amino acid substitutions (M28Q and M68Q) but has a protein yield that is 4 to 5 times higher. We did not observe any ill effects of these substitutions, and this variant was chosen for further analysis.

We determined a crystal structure of EXG:CBM^QQQW^ to enable us to interpret our biochemical experiments more precisely, and to use it as starting point for our molecular dynamics simulations. In the published NMR solution structure of EXG:CBM, the first 3-4 residues were unstructured [10], and so in order to aid crystallization of the new charge-free variant, we produced a truncated version with the four residues deleted from the N-terminal. Our crystals diffracted to 2.2 Å resolution (PDB ID, 6QFS) with detailed data collection and refinement statistics found in the supporting information (Table S1). The eight molecules of EXG:CBM^QQQW^ in the asymmetric unit were conformationally homogeneous and well-superimposable (Figure S1A and C), and displayed close structural similarity to the NMR structure of wild-type EXG:CBM [10] with an aligned backbone RMSD of 1.5 Å. This supports our previous assertion, that removing all charged residues from EXG:CBM had minimal effect on the structure. We have used this crystal structure, which remarkably represents the first in the PDB of a protein of more than 100 amino acids completely free of ionizable amino acid residues, throughout the present work.

### Systematic Insertion of Ionizable Residues

We aimed to analyze the interaction between pairs of charged residues on the surface of EXG:CBM^QQQW^, whose only formal charges before insertion are on the N- and C-termini. To monitor charge interactions, we introduced a histidine (T66H mutation) at the protein surface in the middle of a β-sheet, assuming this would result in minimal conformational changes when introducing other charges (Figure 1A). We then inserted, one at a time, three aspartic acid residues around H66 creating: D39-H66 (T39D mutation), D43-H66 (S43D mutation), and D61-H66 (S61D mutation). The three aspartates were placed with comparable C^α^-C^α^ distances of 7-10 Å (Figure 1A), typical for aspartate-histidine salt bridges [11, 12], but with different degree of solvent exposure (Figure 1B). Neither the N-nor C-terminal charges were expected to interact significantly with this part of the protein. The N-terminus is unstructured, placed on the opposite face of the H66 and did not show any signs of interactions to the rest of the protein in wild-type EXG:CBM [9]. The C-terminus is located around 20 Å away from H66 on the opposite face of the protein and should therefore also have negligible interaction with all the introduced charges.

**Figure 1.**
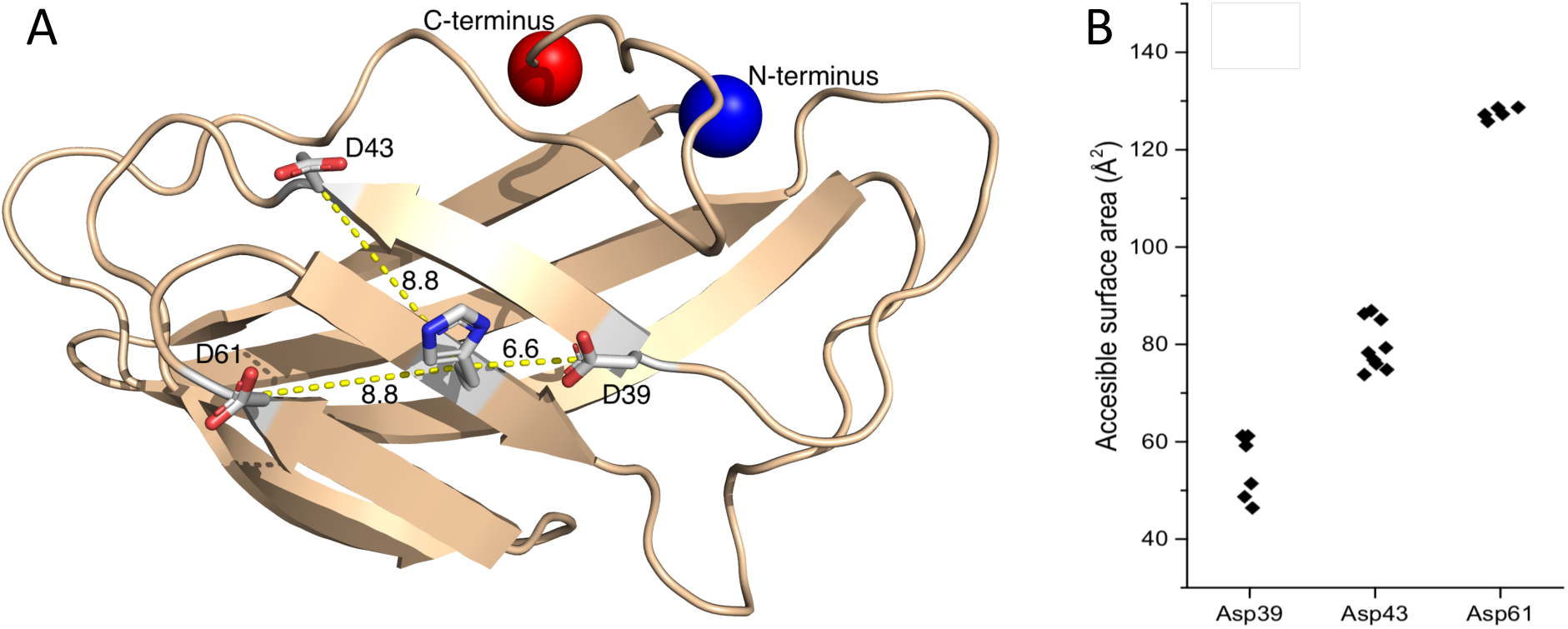
A) Structure of EXG:CBM^QQQW^ with the introduced residues shown in stick representation and the charged N- and C-terminus clearly visualized. These residues are not present in the original crystal structure but are instead introduced here for a representative purpose. Distances between C_α_ atoms of the aspartate residues and the histidine residue is shown. B) Solvent accessible surface area (SASA) of the carboxylic acid group of an aspartate in the three single aspartate containing variants, with points representing the side-chain conformations from the Pymol rotamer library.

### Excellent Agreement in p*K*_a_ Determination by CpHMD and NMR Spectroscopy

We determined the p*K*_a_ values of all the titratable histidine and aspartate residues for all the designed EXG:CBM variants using both CpHMD and NMR spectroscopy (Figure 2 and Table S2, Figure S2-S3). All single and double mutants were characterized, except for the experimental p*K*_*a*_ measurement of the D39. This variant which showed low expression levels which precluded direct measurement by NMR and its p*K*_*a*_ was instead inferred using stability data (see methods). Comparing the p*K*_a_ values predicted from the CpHMD with the values obtained by NMR revealed a very good agreement with a RMSE of 0.39 pH units, which was lower or equal to many model development studies [13] and studies aimed towards protein characterization [14]. With the exception of D39 in the D39-H66 variant, we found a small but systematic overestimation of p*K*_a_ values for the aspartate residues and underestimation for the histidine residues by CpHMD.

**Figure 2.**
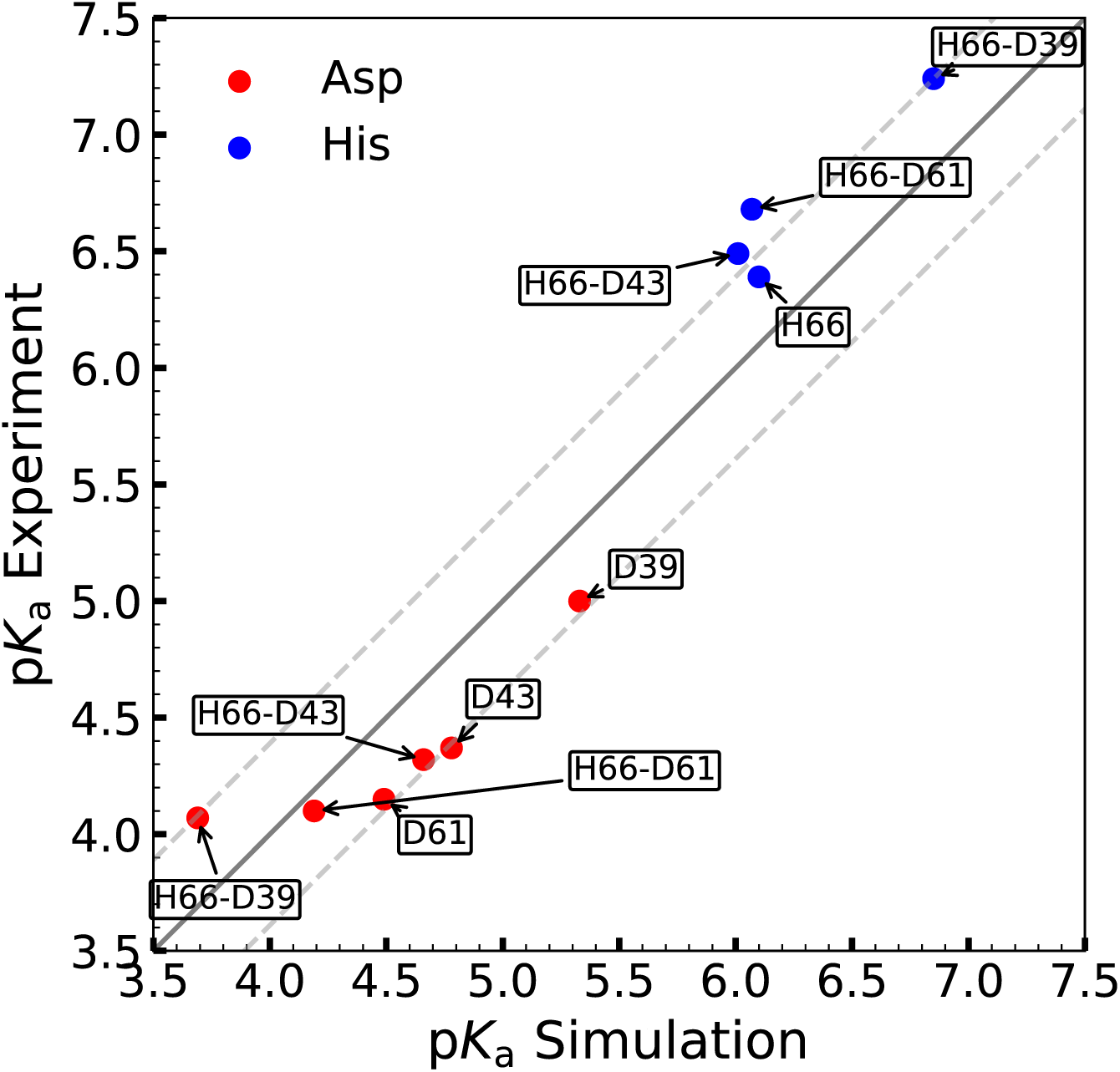
Comparing pK_a_ values from CpHMD with experimental pK_a_ values from NMR spectroscopy. Each point corresponds to a specific residue (red aspartate; blue histidine) in a specific EXG variant (label) so that e.g. the blue point labelled H66-D43 represents the pK_a_ values of H66 in the EXG variant H66-D43.The full line is the diagonal and the dashed lines represent the RMSE (0.36 pK_a_ units) between experiment and computation.

As expected, we found that the presence of H66 shifted the p*K*_*a*_-values of the three Asp residues downwards. Examining the magnitude of this effect we found that while both D43 and D61 were relatively weakly affected by the presence of H66, the p*K*_*a*_-value of D39 was shifted substantially by the presence of H66 (Figure 2). Looking at the experimental p*K*_*a*_-values we found both D43 and D61 were shifted down by 0.05 units, whereas D39 was shifted by approximately 1 unit suggesting a minor interaction. These observations are corroborated by the CpHMD results, thus validating that simulations can be used to interpret the structural origin of the p*K*_a_ shifts. We note, however, also that the effect of H66 appeared to be larger in the simulations compared to the experiments, suggesting that the charge-charge interactions in computation might be overestimated (see below). Turning to H66 we found a similar picture. Specifically, we found that while the p*K*_a_ value of H66 was only mildly affected by D43 and D61, the p*K*_*a*_-value of H66 was upshifted by 0.85 p*K*_a_-units (equivalent to 4.9 kJ/mol), in the presence of D39.

Inspecting the titration profiles obtained from NMR spectroscopy we observed that all the charged residues in the single-charged variants showed single-step titrations with Hill-coefficients estimated to be approximately 1. We also observed monophasic titration profiles from both the D43-H66 and D61-H66 variants, while all the signals from the D39-H66 variant showed biphasic titration profiles (Figure S2). We note that we observe unusually high chemical shifts for the C^γ^ of D39 in this D39-H66 variant, in particular at low pH when the residue becomes protonated. In the simulations at pH 2 we observed in 23% of the structures that the protonated carboxylic acid of D39 acted as a hydrogen bond donor with Q68 being the acceptor, and also observe a minor population of hydrogen bonds from D39 to N103 (3.7%), T41 (0.5%), and H66 (0.4%). While the origin of the unusual chemical shifts of D39 remain unclear we speculate that these hydrogen bond interactions with D39 could play a role in explaining this observation.

In agreement with the NMR spectroscopy data, CpHMD also predicted the titration profile for aspartate and histidine to conform fairly well to the Henderson-Hasselbalch equation for all variants with the exception of the D39-H66 pair, which exhibited biphasic titration curves (Figure S3). The biphasic shape and the symmetry of the D39-H66 titration curves clearly showed that in this variant the aspartate protonation state was dependent on the histidine protonation state and *vice versa*. We used a diprotic acid model, in which the two titratable sites together could adapt four microstates (thus ignoring the histidine tautomerism) [15], to fit the CpHMD data from D39-H66 (Figure 3), and derived a coupling energy between the two non-independent titrations of the D39-H66 pair to be 14.6 kJ/mol. The large coupling energy emphasized that the condition of both the aspartate and histidine being simultaneously protonated or deprotonated was much more energetically unfavorable than the zwitterion and doubly neutral state. Due to strong coupling between D39 and H66, the p*K*_a_ value for the two residues was obtained from the diprotic acid model through macroscopic p*K*_a_ values (figure 2), yielding a p*K*_a_ of 3.69 and 6.85 for the first and second titration event respectively. These two p*K*_a_ values corresponded to the pH of the first and second transition of the biphasic titration curve and should not be confused with single site-specific p*K*_a_ values (known as p*K*_1/2_), which cannot be obtained from the D39-H66 pair. It has previously been shown that the difference between p*K*_1/2_ and macroscopic p*K*_a_ values for diprotic acids is zero when the coupling energy is either zero or very strong [15], thus, we could safely assume the macroscopic p*K*_a_ values obtained from CpHMD was comparable to the experimentally measured p*K*_a_ values for the D39-H66 pair.

**Figure 3.**
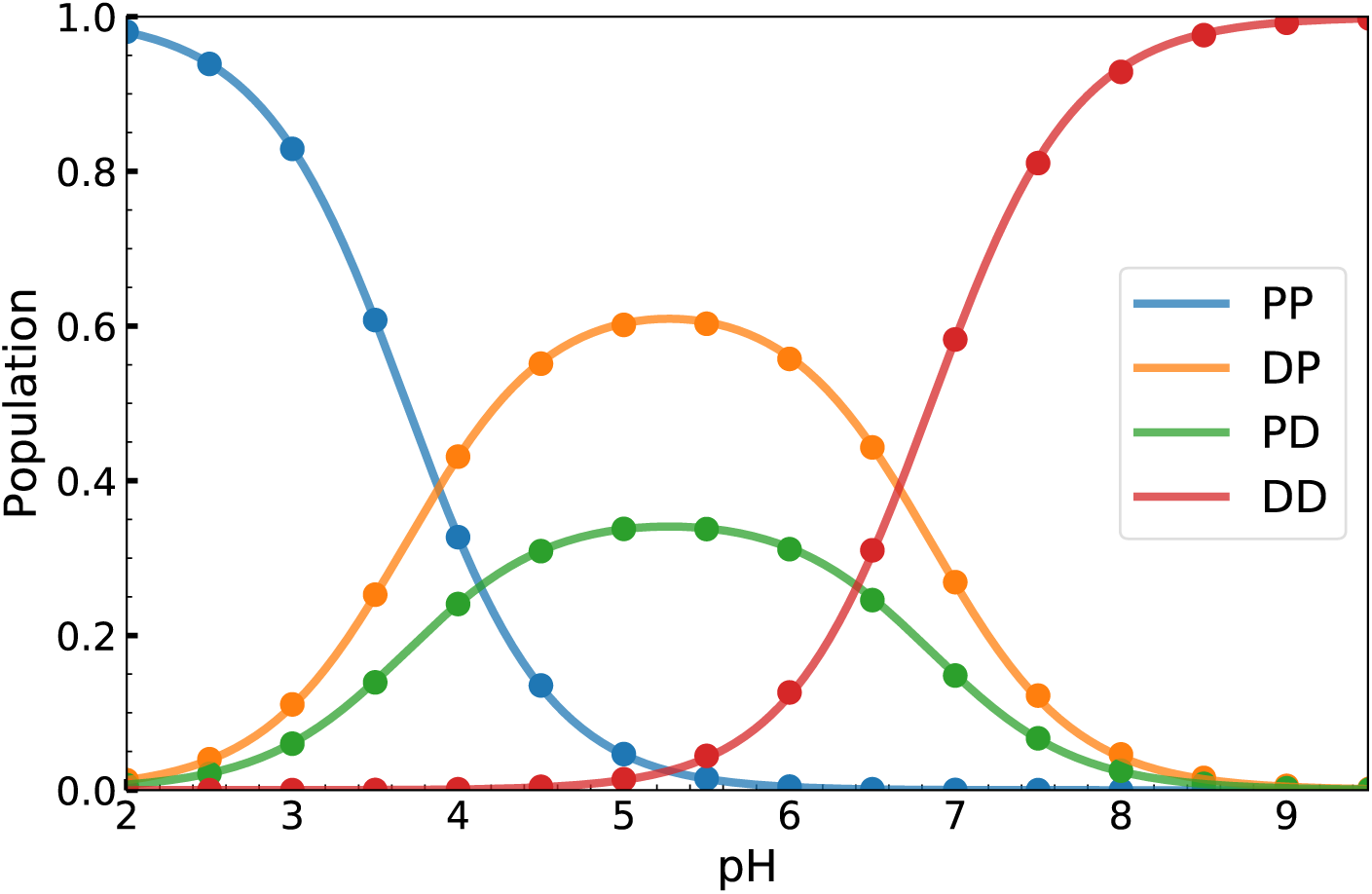
Bjerrum diagram of the diprotic D39-H66 EXG variant with the legend specifying the protonation state of D39 (first) and histidine (second) to be either protonated (P) or deprotonated (D). The points are sampled by CpHMD, while the solid line is the best fit obtained from the diprotic acid model. The microscopic pK_a_ values (where the subscript is the reactant and the superscript is the product) obtained from the fit are 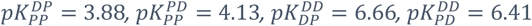, and a coupling energy of 14.6 kJ/mol.

### D39-H66 Pair Adapt Protonation State Dependent Conformation Optimal for Salt Bridging

To investigate the molecular details of the strong interaction of the D39-H66 pair, we used the CpHMD simulations to perform a structural analysis. We calculated the distribution of distances between the N^δ^ and N^ϵ^ atoms in H66 and center of the two side chain oxygens of D39 (Figure 4). At pH 5, we observed a distinct peak around 2.8 Å between the N^ϵ^ and the aspartate carboxylate center of charge. At this pH, the His-Asp system was almost fully in a single-protonated state with ∼60% having a protonated His and negatively charged Asp and ∼40% having a protonated Asp and neutral His a (Figure 2), leading to strong direct interactions between the two residues. Examining the structure corresponding to the short-distance peak we indeed observed that the N^ϵ^ and associated titratable proton of the histidine side chain was facing the carboxylate group (Figure 4). Both the zwitterionic and neutral states possess hydrogen bonding capacity, however, the zwitterionic state additionally possessed formal charges for the aspartate and histidine residue thus causing the interaction to resemble a combination of Coulombic interaction and hydrogen bonding i.e. a salt bridge.

**Figure 4.**
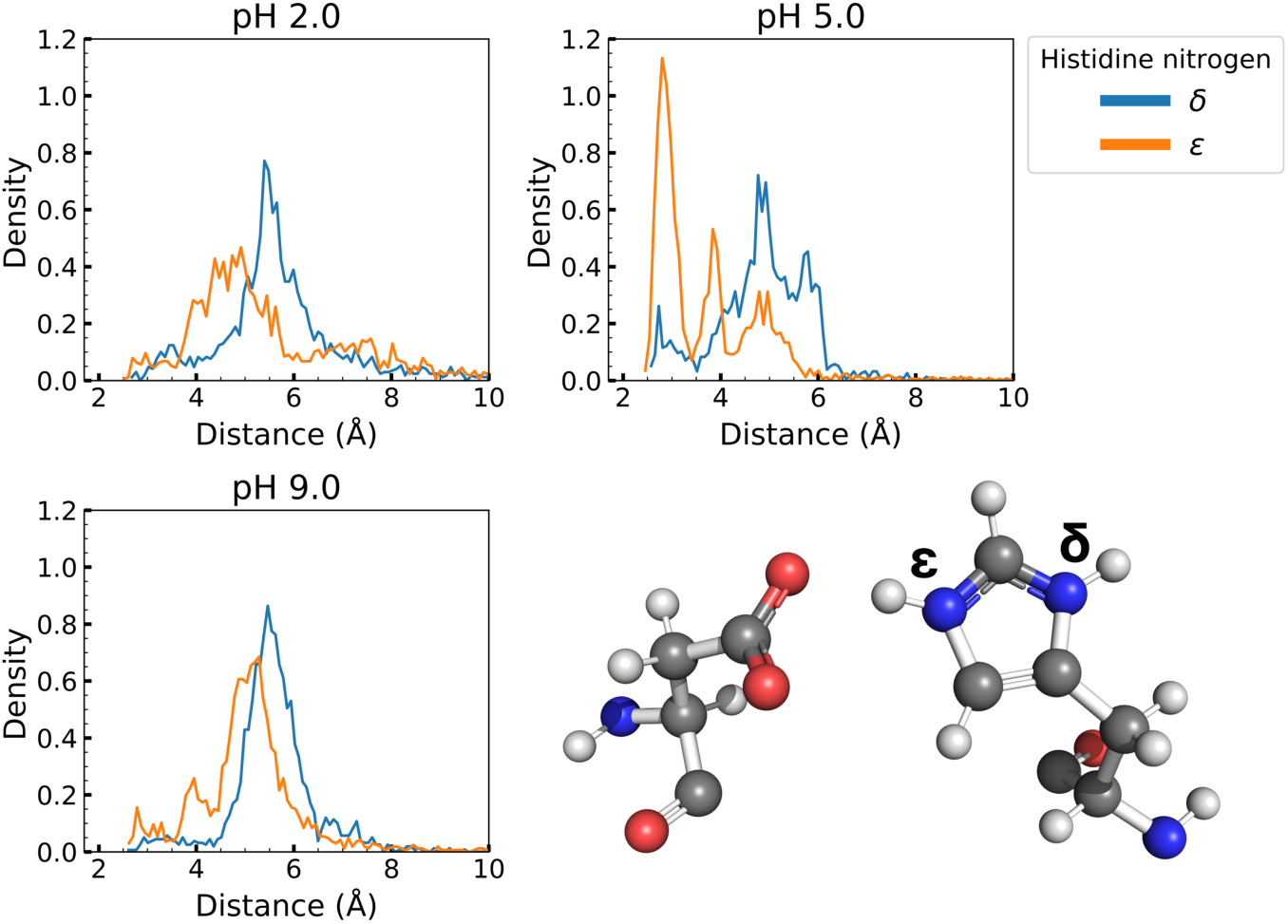
Distance distribution between the N^δ^ or N^ϵ^ (visualized in the bottom right) of His66 and the center of charge of the oxygen in the carboxylate group in the side chain of Asp39 for the Asp39-His66 pair at pH 2.0, 5.0, and 9.0 as obtained from the molecular dynamics simulation.

**Figure 5.**
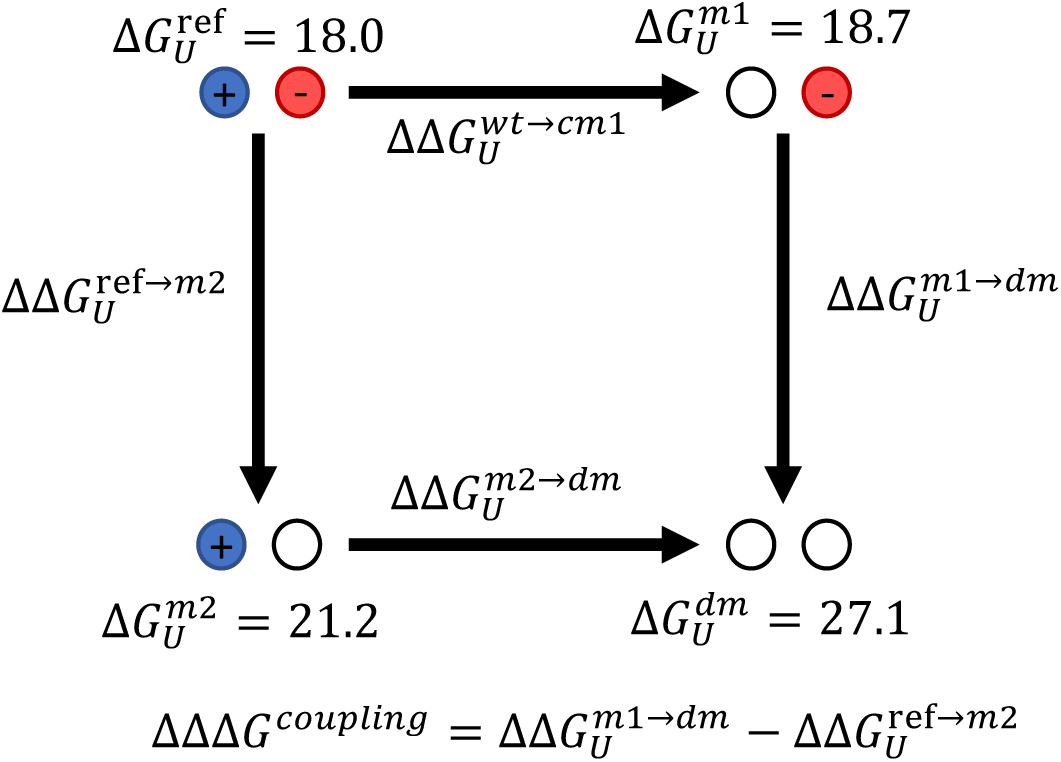
Double-mutant cycle (DMC) relating the unfolding energies, ΔG_U_, of a protein with two charged residues (ref), two single mutants (m1 and m2), and a double mutant with no charged residues (dm). The blue (+), red and (-) white (neutral) balls represent the charge state of the two residues investigated. The ΔG_U_ values shown (in kJ/mol) for the DMC of the D39-H66 variant at pH 8.0 for illustration.

At pH 2.0 and 9.0, where the two residues were either both protonated or deprotonated, respectively, we instead observed a distribution of longer distances to N^ϵ^ between ∼4Å to ∼6Å, that were also found as a minor population at pH 5.0 (Figure 4). Examining the joint distribution of *χ*_1_ angles of D39 and H66, we found that all nine combinations of the three *χ*_1_ rotamers for each residue were sampled, though we saw varying populations that depended substantially on pH (figure S7). At pH 2.0 and 9.0 D39 was mostly found with *χ*_1_ in the **p**-rotamer (*χ*_1_ ∼60°) [16], whereas at pH 5.0 it mostly sampled the **t**-rotamer (*χ*_1_ ∼180°). The histidine was mostly found in the **t**-rotamer, sampling also the **m**-rotamer (*χ*_1_ ∼-60°) at low pH. When D39 was in the **p**-rotamer, the aspartate side chain faced the solvent, whereas the **t**-rotamer state populated at pH 5.0 corresponded to the conformation from the short-distance peak at pH 5 in Figure 4, with the aspartate side chain facing the side chain of H66. These pH-dependent conformations and their exchange can be explained from a simple charge-charge perspective; an unpaired charge will face the solvent to minimize its burial character, while the paired charge of D39 and H66 will face one another to engage in strong interactions, predominantly salt bridging.

The NMR experiments also provided some structural insight into the aspartate-histidine pairs, because the tautomeric state of a protonated histidine could be determined from its C^δ^ chemical shift [43]. In the D39-H66 variant the C^δ^ chemical shift of His66 decreased at the beginning of the H66 titration, consistent with the N^ε^H tautomer being dominant, but at higher pH the signal disappeared (Figure S2). In the other variants, the C^δ^ peak also disappeared at pH coinciding with the histidine titration, suggesting that the variants had a mixture of both tautomers in intermediate exchange. The three His-C^δ^ signals were also affected differently by the protonation of the three different aspartates. Interestingly, they seemed to react differently at the beginning of the H66 titration, indicating that the three aspartates influenced the tautomer distribution differently.

### Partial Burial of Surface-Exposed Ionizable Residues Influence p*K*_a_ Values

The p*K*_a_ values obtained from the titration of the single aspartate variants, by both CpHMD and NMR spectroscopy, differed by up to approximately 1 p*K*_a_ unit despite all being on the protein surface. We attribute the differences in p*K*_a_ values to the difference in partial burial (partial dehydration) of the aspartate residues, which corresponds well with the upshift in p*K*_a_ values and is due to favoring of the carboxylic acid group over the carboxylate anion in a low dielectric environment. As a consequence, the p*K*_a_ of D39 was higher than the p*K*_a_ of D43 which again was higher than that of D61. The p*K*_a_ value of D61 was very close to the reference value for aspartate in the CpHMD model, which could be considered a reference of a fully solvated aspartate. The presence of hydrogen bonds could be another possible contribution to the perturbed p*K*_a_ values. However, from the CpHMD simulations we did not find other residues to engage in long-lived hydrogen bonding with the aspartates, thus supporting the hypothesis that p*K*_a_ differences are due to the different degrees of hydration. These effects appear to be captured by the generalized Born solvation model and corresponds well to SASA data obtained from the crystal structure of EXG:CBM^QQQW^.

The CpHMD simulation also provided some interesting insight into the different dynamics of the three aspartate residues. Being located at the protein surface, the aspartate side chains were free to align along the protein backbone or to stretch out into the solvent. This conformational rearrangement could influence the p*K*_a_ value of a titratable group due to the energetic relief upon a change in protonation state. By looking at the first dihedral angle of the aspartate side chain (χ_1_), this pH dependent conformational change could be observed for the D39 variant, while it was much less pronounced in the D43 and D61 variants (Figure S8). The absence of long-lived hydrogen bonds between the charged residues and the backbone protons, and the fact that no significant structural rearrangements take place in the simulations, also agrees with the experimental observations. This is also in agreement with the observation that the H^N^ chemical shifts in all the EXG:CBM variants (except for the H^N^-shifts of the charged residues) changed with less than 0.2 ppm in response to the pH titration.

### Thermodynamic Cycles Reveals Different Modes of His-Asp Interaction

The results described above demonstrated substantial perturbation of the p*K*_*a*_-values of several of the residues, and also highlighted strong coupling between the titration behavior of Asp39 and His66. To get an alternative view of the coupling between the residues in the three charge pairs, we turned to thermodynamic double mutant cycles (DMC). In DMC, the change of the stability from removing a charged residue in a charge pair was subtracted from the change of the stability from removing the same residue in a protein variant lacking the charged partner (Figure 4). In our analysis, we treated the charge pair as the reference state, and mutated the protein sequence back to the original EXG:CBM^QQQW^ sequence. We performed the DMC experiments at several different values of pH and also in some cases in the presence of high concentrations of salt.

All the tested variants displayed classical two-state equilibrium unfolding behavior (Figure S4, for individual unfolding free energies see Table S3). Also, comparing mutant variants, only very few changes in the chemical shifts were observed in ^15^N-HSQC NMR spectra, suggesting the structures to be very similar (Figure S5). The stability data and thermodynamic cycles are presented in Figure S4 and Figure S6, and the summarized results of apparent coupling energies are shown in Table 1. The DMC data confirmed that, although all three aspartates are placed at potential salt bridge distances to the histidine, their interactions were very different.

**Table 1.**
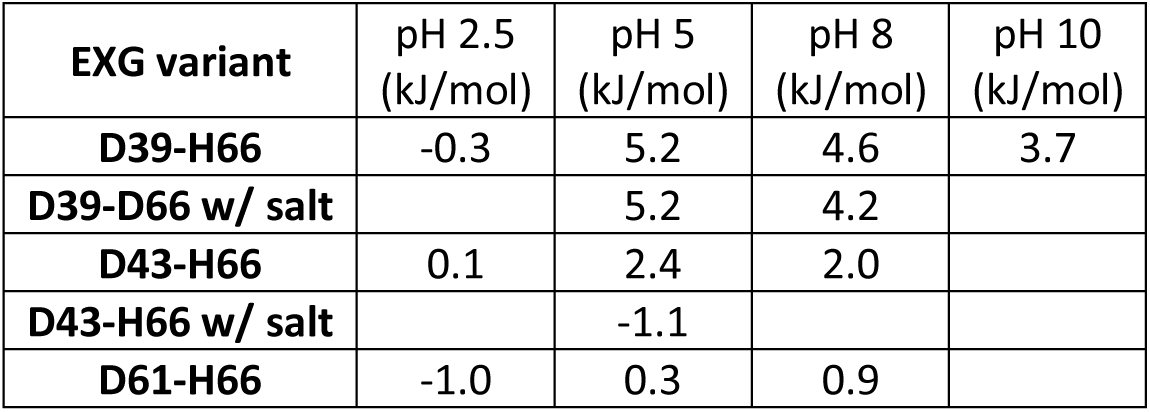
Apparent coupling energies from double-mutant cycles

| EXG variant | pH 2.5<br>(kJ/mol) | pH 5<br>(kJ/mol) | pH 8<br>(kJ/mol) | pH 10<br>(kJ/mol) |
| --- | --- | --- | --- | --- |
| D39-H66 | -0.3 | 5.2 | 4.6 | 3.7 |
| D39-D66 w/ salt |  | 5.2 | 4.2 |  |
| D43-H66 | 0.1 | 2.4 | 2.0 |  |
| D43-H66 w/ salt |  | -1.1 |  |  |
| D61-H66 | -1.0 | 0.3 | 0.9 |  |

We focus first on the results at pH 5, where the His and Asp residues will generally have taken up a single proton, generally associated with H66. In line with the strong coupling of the titration behavior of D39-H66, we found that coupling energy from the DMC was also higher for this pair than the other two (Table 1). Protonating the aspartates at pH 2.5 decreased the coupling energies to close to zero, suggesting that the charge interactions between the Asp and His pairs contributed substantially to the coupling energies measured at pH 5.

Increasing pH to 8.0 or adding salt, however, revealed a more complex picture. While the couplings generally become smaller in these cases, they remained large, particularly for D39-H66. At pH 5.0 the coupling free energy of D39-H66 remained constant at 5.2 kJ/mol even after addition of 1.5 M NaCl and only dropped to 4.6 kJ/mol and 3.7 kJ/mol at pH 8 and 10 respectively. Thus, these results suggest a strong interaction between D39-H66 relative to the reference state, even when both residues were deprotonated.

Overall, the data therefore suggest three different cases in the three aspartate-histidine pairs. D61 does not seem to interact with H66 to a measurable degree with any of the approaches used in this study. The interaction between D43 and H66 is relatively week and can be screened by salt, pointing at it being classical Coulombic in nature. The reason why this interaction is not more pronounced in the NMR and CpHMD data, could perhaps be the higher salt concentrations (0.1 M NaCl) in these experiments. The strong interaction between D39 and H66 can, on the other hand, not be explained purely by a Coulombic interaction since the interaction persist at pH 10 where H66 is not positively charged and cannot be screened by 1.5 M salt. The latter is, however, not unusual for solvated [17] and partially solvated [18] salt bridges.

## Concluding Remarks

We have designed and explored a simple model system to study protein electrostatics, looking at three different positions for Asp residues surrounding a His residue. We found differences in hydration, even for residues that one would typically categorize as solvated or partially solvated, which yielded a difference in p*K*_a_ up to approximately 1 unit, equivalent to approximately 5.7 kJ/mol. However, the perturbation by partial burial was only secondary compared to the impact of the interaction between the histidine and aspartate residues. From the three Asp-His pairs studied here, only one engaged in a salt bridge, causing strongly coupled titration curves. Thus, CpHMD simulations suggested the interaction between the two strongly coupled titration sites to favor salt-bridging over hydrogen bonding obtained (60:40) from the proton tautomerization. This was most likely due to the charge-pair being partially solvated on the surface of EXG. Finally, due to the generally good agreement between the CpHMD simulations and NMR experiments, we were able to gain detailed mechanistic insight into the relationship between protein interactions and titrations. Specifically, the biphasic titration observed here was caused by strong electrostatic interactions and the observed pH-dependent coupling energies became easier to interpret. As of such, we anticipate this work will expand the general knowledge of protein electrostatics studies.

## Materials & Methods

**Molecular dynamics simulations** were performed in the Amber16 PMEMD (Particle Mesh Ewald Molecular Dynamics) program [19,20] with long range electrostatics handled by particle mesh Ewald summation [21] and default LJ and short-range electrostatic cutoffs, using the AMBER constph force field, derived from ff14SB [22], in explicit TIP3P water [23]. All simulations were run with pH replica exchange enhanced sampling [24,25] in the pH range 2.0-9.5 for the double titratable variants and single histidine variant, and pH range 0.5-8.0 for the single aspartate variants with all replica interspaced by 0.5 pH units with each replica running 100 ns. The systems were first minimized and next equilibrated in terms of temperature and pressure to 300 Kelvin and 1 bar pressure separately by Langevin dynamics [26,27] with a collision frequency of 5.0 ps^-1^ and the Berendsen barostat [28,29] with a rise time (coupling constant) of 1 ps and isothermal compressibility of 44.6 × 10^−6^ bar^- 1^. A time step of 2 fs was utilized and SHAKE algorithm [30] applied to constrain hydrogen-heavy atom bond lengths. The equilibration was followed by a production run in the semi-grand canonical ensemble of protons (*NVΔμ*_*H*_^*+*^) where, only histidine and aspartate residues were allowed to titrate every 0.2 ps through a Monte Carlo protonation state move for each titratable residue and every 0.4 ps attempt to perform a replica exchange between the simulations to swap pH. The Monte Carlo protonation state move used Generalized Born solvation [13,31] with a salt concentration (Debye-Hückel based) of 0.1 M monovalent salt [32] and upon a successful change in protonation state 0.2 ps of solvent relaxation was conducted after reinstating the explicit water [13]. Configurations were gathered every 50 ps for statistical evaluation.

For single-phasic titration curves, p*K*_a_ values and Hill coefficients were obtained through averaging the deprotonated factions for each pH of the simulation, fitted to the Hill modified Henderson-Hasselbalch equation [33-35] yielding a single p*K*_a_ value and Hill coefficient for each titratable residue. This method is very much equivalent to titration experiments and thus effectively measure the ensemble averaged deprotonated fractions. In these analyses we included a Hill coefficient as a control to see whether the transitions would be broadened, but found n_H_∼1 in all cases. For biphasic titration curves, the population of the protonation species for each pH were fitted to a diprotic acid model, yielding macroscopic and microscopic p*K*_a_ values as explained in detail by Ullmann [15].

**Protein expression and purification** of all variants were done following the same protocol as described for EXG:CBM [9], although the gel filtration step and second lyophilization were omitted and the temperature needed to elute the protein from Avicel (Sigma-Aldrich) ranged from 50 to 70 °C. The protein yield after elution from Avicel were ∼30 mg/L E. coli culture for EXG:CBM^QQQW^ and EXG:CBM^QQQW,66H^, 11-18 mg/L for EXG:CBM^QQQW,43D^, EXG:CBM^QQQW,61D^, EXG:CBM^QQQW,39D,66H^, EXG:CBM^QQQW,43D,66H^, and EXG:CBM^QQQW,61D,66H^, and 3-6 mg/L EXG:CBM^QQQW,39D^. After purification, the proteins were buffer exchanged into the appropriate buffer using an Illustra NAP-5 column (GE Healthcare).

**Protein NMR** were performed on a Bruker 600 MHz Avance IIIHD spectrometer with a quadruple resonance cryoprobe. The amide chemical shifts of EXG:CBM^QQQW,39D,66H^ were assigned from NH(CA)CO, HNCO, HNCOCACB and HNCACB spectra at pH 3.6. Signals for all residues except Ala1 and Cys108 could be identified. The combined assignment of EXG:CBM^QQQW,39D,66H^ and EXG:CBM [9] made it possible to assign the amide signals in the remaining variants. To get the C^ϵ1^, H^ϵ1^, C^δ2^, and H^δ2^ chemical shifts of His, spectra of the aromatic carbon region was measured (using the Bruker pulse sequence *trosyargpphwg*), and a H2(C)CO experiment, correlating C^γ^ chemical shifts to H^β^, was measured in order to get the p*K*_a_ of the aspartates (Bruker pulse sequence *hcacogp3d* optimized for carboxylic side chains [36]). Because the C^α^ and the carboxylic acid of the C-terminal glycine resembles an aspartate side chain, the chemical shifts of the C-terminal was also directly measured in the H2(C)CO experiment. Side chain assignment of the relevant residues was easy, since each spectrum only contained one set of Asp, His or C-terminal signals. For the titration, NMR samples (0.3 mM, 100 mM KCl, 10% D_2_O, 1 mM DDS) were split in two (one titrating up, one titrating down), and the appropriate side chain spectra together with a N^15^-HSQC were measured in steps of 1/3 pH units at 25 °C. Titration curves were fitted to the modified Hill equation [33-35], while chemical shifts with two titration events were fitted to equation (eq. 1):

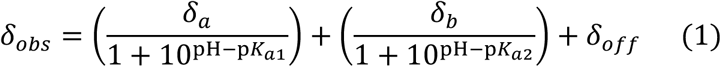

Where δ_*obs*_ is the observed chemical shift, δ_*a*_ and δ_*b*_ are the change in chemical shift due to the first and second titration with *pK*_*a*1_ and *pK*_*a*2_ respectively, and δ_*off*_ is the offset. For fitting, the chemical shifts of C^γ^ (Asp), C^ϵ1^ (His) or C (C-term) were used. In case of incomplete titrations of the side chain signals, the amide and side chain ^13^C signal of a changed residue were globally fit to obtain the pK_a_ value. The pK_a_ value of EXG:CBM^QQQW,66H^ were fitted from amide signals.

### Protein stability and double mutant cycles

All experiments were conducted in 5 mM buffer (pH 2.5, sodium phosphate; pH 5, sodium acetate; pH 8, MOPS; pH 10, boric acid) Urea solutions were made fresh daily and incubated with AG 501-X8 resin (Biorad) for minimum 5 hours to remove ionic contaminants. The refractive index was measured after the addition of buffer compound in order to determine the exact concentration of each urea stock. Urea, buffer and protein were mixed to the appropriate concentrations and left to equilibrate at room temperature overnight before measurements. The fluorescence of each sample was measured on a PerkinElmer LS 55 at 25 °C. Due to relatively weak transitions between folded and unfolded state, fits to the linear extrapolation method [37] was made globally for six fluorescence signals of each sample (Em.: 280 and 295 nm, Ex.: 360, 370, and 380 nm) to obtain C_m_-values and apparent m-values, m_app_. ΔG-values were calculated from the linear extrapolation model [37] [*C*_*m*_ · ⟨*m*_app_⟩, ⟨*m*_app_⟩ being the average of *m*_app_ from all measurements. Apparent coupling energies (Figure S9) were calculated as the difference between opposite sites in a thermodynamic cycle (i.e.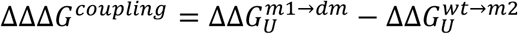). The error on individual stability determinations was in the order of 0.7 kJ/mol, based on four independent experiments on the same variant (D39-H66). With error propagation, this amounts to approximately 1.4 kJ/mol for the coupling energies. Estimation of error is, however, somewhat complicated by the global averaging of m-values which may affect some stability determinations more than others.

For the D39 variant, a p*K*_*a*_ value by NMR titration experiments could not be determined due to low expressions levels, however the p*K*_*a*_ has instead been estimated from unfolding data. The p*K*_*a*_ of the folded state of D39 can be estimated to be 1.1 p*K*_*a*_-unit higher than in the unfolded state. Assuming that the value in the unfolded state is close to the reference value of aspartate, the p*K*_*a*_ of D39 would be upshifted to ∼5.

### Protein crystallization and structure determination by X-ray diffraction

A truncated EXG:CBM^QQQW^ variant missing the N-terminal residues 2-5 was crystallized by vapor diffusion hanging drop experiment. EXG:CBM^QQQW^ at a concentration of 32 mg/mL was mixed at a 1:2 ratio with the crystallization precipitant solution (450 mM MES monohydrate pH 6.5, 55 mM HEPES pH 7.5, 45 mM ZnSO_4_ heptahydrate, 27.5 mM CdSO_4_ hydrate, 550 mM Na acetate trihydrate, 11.25% PEG monomethyl ether 550) and incubated at 10°C. Crystals appeared within a week, and were cryoprotected prior to cryostorage by mixing 1:1 with a cryoprotectant of 28% ethylene glycol, 19.3% sucrose, 120 μg/mL 3-(1-Pyridinio)-1-propanesulfonate. X-ray diffraction data were collected on a PILATUS 6M detector (Dectris) at the ESRF beamline ID30B at 100 K and a wavelength of 0.954 Å.

Diffraction data were indexed and integrated in XDS [38], scaled and merged using Aimless [39], and phases solved by molecular replacement in Phaser [40] with the NMR structure of EXG:CBM (PDB ID, 1EXG) as search model [10]. Maximum likelihood refinement was performed in *phenix*.*refine* [41] against data to a maximum resolution of 2.2 Å with manual model correction in COOT [42]. No disallowed peptide torsion angles were modeled, with 95.64% Ramachandran favored and no outliers. Data collection and refinement statistics are summarized in Table S1. The final coordinates have been deposited in the PDB with the accession code 6QFS.

## ACKNOWLEDGEMENTS

This work was supported by a grant from the Velux Foundations. SHH thanks LUNARC in Lund, Sweden for computational resources.

## Author contributions

SHH, CH, KEJ, KLL, and JRW designed research; SHH, CH, KEJ, YW, KW, and DY performed research; SHH, CH, YW, JM, KT, KLL, and JRW analyzed data; and SHH, CH, KEJ, KLL, and JRW wrote the paper.

The authors declare no competing interest.

The experimental NMR data are electronically available at http://doi.org/10.5281/zenodo.397915 Scripts, data and analysis for the simulations are electronically available at https://github.com/KULL-Centre/papers/tree/master/2020/EXG-charges-hervoe-hansen-et-al

## Notes

### Competing Interest Statement

The authors have declared no competing interest.

### Summary of Updates

We have included links to on-line data.

## References

1. A. Warshel, et al., Electrostatic Basis for Enzyme Catalysis. Chemical Reviews 106, 3210–3235 (2006).

2. R. Varadarajan, T. Zewert, H. Gray, S. Boxer, Effects of buried ionizable amino acids on the reduction potential of recombinant myoglobin. Science 243, 69–72 (1989).

3. H. Luecke, Proton Transfer Pathways in Bacteriorhodopsin at 2.3 Angstrom Resolution. Science 280, 1934–1937 (1998).

4. D. A. Doyle, et al., The Structure of the Potassium Channel: Molecular Basis of K+ Conduction and Selectivity. Science 280, 69–77 (1998).

5. S. Xiao, et al., Rational modification of protein stability by targeting surface sites leads to complicated results. Proceedings of the National Academy of Sciences 110, 11337–11342 (2013).

6. D. G. Isom, C. A. Castaneda, B. R. Cannon, B. Garcia-Moreno E., Large shifts in pKa values of lysine residues buried inside a protein. Proceedings of the National Academy of Sciences 108, 5260–5265 (2011).

7. R. L. Thurlkill, G. R. Grimsley, J. M. Scholtz, C. N. Pace, Hydrogen Bonding Markedly Reduces the pK of Buried Carboxyl Groups in Proteins. Journal of Molecular Biology 362, 594–604 (2006).

8. J.-H. Cho, D. P. Raleigh, Mutational Analysis Demonstrates that Specific Electrostatic Interactions can Play a Key Role in the Denatured State Ensemble of Proteins. Journal of Molecular Biology 353, 174–185 (2005).

9. C. Højgaard, et al., A Soluble, Folded Protein without Charged Amino Acid Residues. Biochemistry 55, 3949–3956 (2016).

10. G.-Y. Xu, et al., Solution Structure of a Cellulose-Binding Domain from Cellulomonas fimi by Nuclear Magnetic Resonance Spectroscopy. Biochemistry 34, 6993–7009 (1995).

11. S. Kumar, R. Nussinov, Close-Range Electrostatic Interactions in Proteins. ChemBioChem 3, 604 (2002).

12. J. E. Donald, D. W. Kulp, W. F. DeGrado, Salt bridges: Geometrically specific, designable interactions. Proteins: Structure, Function, and Bioinformatics 79, 898–915 (2011).

13. J. M. Swails, D. M. York, A. E. Roitberg, Constant pH Replica Exchange Molecular Dynamics in Explicit Solvent Using Discrete Protonation States: Implementation, Testing, and Validation. Journal of Chemical Theory and Computation 10, 1341–1352 (2014).

14. F. Hofer, V. Dietrich, A. S. Kamenik, M. Tollinger, K. R. Liedl, pH-Dependent Protonation of the Phl p 6 Pollen Allergen Studied by NMR and cpH-aMD. Journal of Chemical Theory and Computation 15, 5716–5726 (2019).

15. G. M. Ullmann, Relations between Protonation Constants and Titration Curves in Polyprotic Acids: A Critical View. The Journal of Physical Chemistry B 107, 1263–1271 (2003).

16. S. C. Lovell, J. M. Word, J. S. Richardson, D. C. Richardson, The penultimate rotamer library. Proteins: Structure, Function, and Genetics 40, 389–408 (2000).

17. S. Pylaeva, M. Brehm, D. Sebastiani, Salt Bridge in Aqueous Solution: Strong Structural Motifs but Weak Enthalpic Effect. Scientific Reports 8 (2018).

18. D. L. Luisi, et al., Surface Salt Bridges, Double-Mutant Cycles, and Protein Stability: An Experimental and Computational Analysis of the Interaction of the Asp 23 Side Chain with the N-Terminus of the N-Terminal Domain of the Ribosomal Protein L9. Biochemistry 42, 7050–7060 (2003).

19. R. Salomon-Ferrer, A. W. Götz, D. Poole, S. Le Grand, R. C. Walker, Routine Microsecond Molecular Dynamics Simulations with AMBER on GPUs. 2. Explicit Solvent Particle Mesh Ewald. Journal of Chemical Theory and Computation 9, 3878–3888 (2013).

20. S. Le Grand, A. W. Götz, R. C. Walker, SPFP: Speed without compromise—A mixed precision model for GPU accelerated molecular dynamics simulations. Computer Physics Communications 184, 374–380 (2013).

21. T. Darden, D. York, L. Pedersen, Particle mesh Ewald: An N-log(N) method for Ewald sums in large systems. The Journal of Chemical Physics 98, 10089–10092 (1993).

22. J. A. Maier, et al., ff14SB: Improving the Accuracy of Protein Side Chain and Backbone Parameters from ff99SB. Journal of Chemical Theory and Computation 11, 3696–3713 (2015).

23. W. L. Jorgensen, J. Chandrasekhar, J. D. Madura, R. W. Impey, M. L. Klein, Comparison of simple potential functions for simulating liquid water. The Journal of Chemical Physics 79, 926–935 (1983).

24. S. G. Itoh, A. Damjanovic, B. R. Brooks, pH replica-exchange method based on discrete protonation states. Proteins: Structure, Function, and Bioinformatics 79, 3420–3436 (2011).

25. J. M. Swails, A. E. Roitberg, Enhancing Conformation and Protonation State Sampling of Hen Egg White Lysozyme Using pH Replica Exchange Molecular Dynamics. Journal of Chemical Theory and Computation 8, 4393–4404 (2012).

26. R. W. Pastor, B. R. Brooks, A. Szabo, An analysis of the accuracy of Langevin and molecular dynamics algorithms. Molecular Physics 65, 1409–1419 (1988).

27. J. A. Izaguirre, D. P. Catarello, J. M. Wozniak, R. D. Skeel, Langevin stabilization of molecular dynamics. The Journal of Chemical Physics 114, 2090–2098 (2001).

28. H. J. C. Berendsen, J. P. M. Postma, W. F. van Gunsteren, A. DiNola, J. R. Haak, Molecular dynamics with coupling to an external bath. The Journal of Chemical Physics 81, 3684–3690 (1984).

29. T. Morishita, Fluctuation formulas in molecular-dynamics simulations with the weak coupling heat bath. The Journal of Chemical Physics 113, 2976–2982 (2000).

30. J.-P. Ryckaert, G. Ciccotti, H. J. C. Berendsen, Numerical integration of the cartesian equations of motion of a system with constraints: molecular dynamics of n-alkanes. Journal of Computational Physics 23, 327–341 (1977).

31. J. Mongan, D. A. Case, J. A. McCammon, Constant pH molecular dynamics in generalized Born implicit solvent. Journal of Computational Chemistry 25, 2038–2048 (2004).

32. J. Srinivasan, M. W. Trevathan, P. Beroza, D. A. Case, Application of a pairwise generalized Born model to proteins and nucleic acids: inclusion of salt effects. Theoretical Chemistry Accounts: Theory, Computation, and Modeling (Theoretica Chimica Acta) 101, 426–434 (1999).

33. L. J. Henderson, Concerning the relationship between the strength of acids and their capacity to preserve neutrality. American Journal of Physiology-Legacy Content 21, 173–179 (1908).

34. T. Hill, Cooperativity theory in biochemistry, steady-state and equilibrium systems. New York: Springer-Verlag (1985).

35. J. L. Markley, Observation of histidine residues in proteins by nuclear magnetic resonance spectroscopy. Accounts of Chemical Research 8, 70–80 (1975).

36. L. E. Kay, M. Ikura, R. Tschudin, A. Bax, Three-dimensional triple-resonance NMR spectroscopy of isotopically enriched proteins. Journal of Magnetic Resonance (1969) 89, 496–514 (1990).

37. R. F. Greene, C. N. Pace, Urea and guanidine hydrochloride denaturation of ribonuclease, lysozyme, α-chymotrpsin, and β-lactoglobumin. Journal of Biological Chemistry 249, 5388–5394 (1974).

38. W. Kabsch, XDS. Acta Crystallographica Section D Biological Crystallography 66, 125–132 (2010).

39. P. R. Evans, G. N. Murshudov, How good are my data and what is the resolution? Acta Crystallographica Section D Biological Crystallography 69, 1204–1214 (2013).

40. A. J. McCoy, et al., Phasercrystallographic software. Journal of Applied Crystallography 40, 658–674 (2007).

41. P. D. Adams, et al., PHENIX: a comprehensive Python-based system for macromolecular structure solution. Acta Crystallographica Section D Biological Crystallography 66, 213–221 (2010).

42. P. Emsley, K. Cowtan, Coot: model-building tools for molecular graphics. Acta Crystallographica Section D Biological Crystallography 60, 2126–2132 (2004).

