## Supplementary material for "Charge Interactions in a Highly Charge-depleted Protein": Manuscript_Sup_v13c.docx

^a^Linderstrøm-Lang Centre of Protein Science, Department of Biology, University of Copenhagen, DK-2200 Copenhagen N, Denmark, ^b^Division of Theoretical Chemistry, Department of Chemistry, Lund University, SE 221 00 Lund, Sweden, ^c^VIB-VUB Center for Structural Biology, Vlaams Instituut voor Biotechnologie - Vrije Universiteit Brussel, B-1050 Brussels, Belgium, ^d^Brussels Center for Redox Biology, Vrije Universiteit Brussel, B-1050 Brussels, Belgium, ^e^Structural Biology Brussels, Vrije Universiteit Brussel, B-1050 Brussels, Belgium

^1^ These authors contributed equally to this work.

^§^ Current address: The Francis Crick Institute, London, UK.

Contents:

Data collection and refeinment statistics (Table S1)

p*K*_a_ values and Hill parameters from titrations of EXG:CBM variants (Table S2)

Stability parameters from urea unfolding (Table S3)

Crystal structure of EXG:CBM^QQQW,Δ2-5^ (Figure S1)

pH titration curves followed by NMR chemical shifts (Figure S2)

pH titration curves followed by CpHMD (Figure S3)

Urea induced unfolding data (Figure S4)

^15^N-HSQCs of the investigated EXG:CBM variants (Figure S5)

Double-mutant cycles (Figure S6)

Rotamer population of Asp and His in double titratable EXG:CBM variants (Figure S7)

Rotamer population of Asp in single titratable EXG:CBM variants (Figure S8)

Thermodynamic cycle for estimating pK_a_ shifts from stability data (Figure S9)

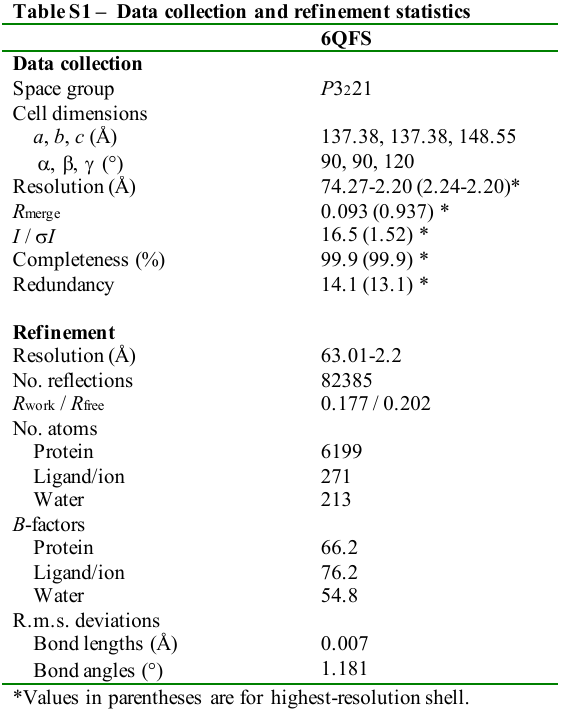

**Table S2 – p*K*_a_ values and Hill coefficients from titrations of EXG:CBM variants.**

Hill coefficients from CpHMD simulation and experimental determination.

|  | Asp39 | Asp43 | Asp61 | His66 |
| --- | --- | --- | --- | --- |
| CpHMD/Experimental values | | | | |
| EXG:CBM^QQQW,66H^ |  |  |  | 1.0/0.98 |
| EXG:CBM^QQQW,39D^ | 1.01/- |  |  |  |
| EXG:CBM^QQQW,43D^ |  | 1.0/1.08 |  |  |
| EXG:CBM^QQQW,61D^ |  |  | 0.96/0.98 |  |
| EXG:CBM^QQQW,39D,66H^ | -/- |  |  | -/- |
| EXG:CBM^QQQW,43D,66H^ |  | 0.97/- |  | 0.98/ 0.91 |
| EXG:CBM^QQQW,61D,66H^ |  |  | 0.94/- | 0.99/1.1 |

^-^ Hill coefficient not determined due to double titration in signal.

pK_a_-values from CpHMD simulation and experimental determination.

|  | Asp39 | Asp43 | Asp61 | His66 |
| --- | --- | --- | --- | --- |
| CpHMD/Experimental values | | | | |
| EXG:CBM^QQQW,66H^ |  |  |  | 6.1/6.39 |
| EXG:CBM^QQQW,39D^ | 5.33/~5^b^ |  |  |  |
| EXG:CBM^QQQW,43D^ |  | 4.78/4.37 |  |  |
| EXG:CBM^QQQW,61D^ |  |  | 4.49/4.15 |  |
| EXG:CBM^QQQW,39D,66H^ | 3.69^a^/4.07 |  |  | 6.85^a^/7.24 |
| EXG:CBM^QQQW,43D,66H^ |  | 4.66/4.32 |  | 6.01/6.49 |
| EXG:CBM^QQQW,61D,66H^ |  |  | 4.19/4.1 | 6.07/6.68 |

^a^Macroscopic p*K*_a_ from the diprotic acid model (see Materials and Methods)

^b^This pK_a_-value is an estimate based on stability data (See Figure S6).

**Table S3 – Stability parameters from urea unfolding**

|  |  | m_app_ (kJ/mol)^a^ | C_m_ (M)^a^ | ΔG_U_ (kJ/mol)^b^ |
| --- | --- | --- | --- | --- |
| EXG:CBM^QQQW^ | pH 2.5 | 4,24 | 5,01 | 26,4 |
|  | pH 5 | 4,79 | 5,16 | 27,1 |
|  | pH 8 | 5,22 | 4,95 | 26,0 |
|  | pH 10 | 5,17 | 4,89 | 25,7 |
|  | pH 5, 1.5M NaCl | 4,47 | 6,33 | 33,3 |
|  | pH 8, 1.5M NaCl | 4,64 | 6,43 | 33,8 |
| EXG:CBM^QQQW,66H^ | pH 2.5 | 3,91 | 4,16 | 21,9 |
|  | pH 5 | 4,51 | 4,04 | 21,2 |
|  | pH 8 | 4,38 | 3,96 | 20,8 |
|  | pH 10 | 4,53 | 3,91 | 20,6 |
|  | pH 5, 1.5M NaCl | 4,77 | 5,56 | 29,2 |
|  | pH 8, 1.5M NaCl | 7,46 | 5,64 | 29,7 |
| EXG:CBM^QQQW,39D^ | pH 2.5 | 5,58 | 4,15 | 21,9 |
|  | pH 5 | 4,95 | 3,56 | 18,7 |
|  | pH 8 | 6,20 | 2,85 | 15,0 |
|  | pH 10 | 6,75 | 2,77 | 14,6 |
|  | pH 5, 1.5M NaCl | 5,13 | 4,47 | 23,5 |
|  | pH 8, 1.5M NaCl | 4,87 | 4,20 | 22,1 |
| EXG:CBM^QQQW,43D^ | pH 2.5 | 5,54 | 4,81 | 25,3 |
|  | pH 5 | 5,06 | 4,66 | 24,5 |
|  | pH 8 | 5,38 | 4,54 | 23,9 |
|  | pH 5, 1.5M NaCl | 4,37 | 6,26 | 32,9 |
| EXG:CBM^QQQW,61D^ | pH 2.5 | 5,40 | 5,11 | 26,9 |
|  | pH 5 | 5,16 | 4,96 | 26,1 |
|  | pH 8 | 4,77 | 4,83 | 25,4 |
| EXG:CBM^QQQW,66H,39D^ | pH 2.5 | 5,21 | 3,25 | 17,1 |
|  | pH 5 | 5,67 | 3,42 | 18.0 |
|  | pH 8 | 5,87 | 2,75 | 14,4 |
|  | pH 10 | 5,77 | 2,50 | 13,1 |
|  | pH 5, 1.5M NaCl | 5,22 | 4,69 | 24,6 |
|  | pH 8, 1.5M NaCl | 4,35 | 4,23 | 22,2 |
| EXG:CBM^QQQW,66H,43D^ | pH 2.5 | 6,35 | 3,97 | 20.9 |
|  | pH 5 | 5,43 | 3,99 | 21.0 |
|  | pH 8 | 5,43 | 3,93 | 20,7 |
|  | pH 5, 1.5M NaCl | 4,20 | 5,26 | 27,7 |
| EXG:CBM^QQQW,66H,61D^ | pH 2.5 | 5,32 | 4,06 | 21,4 |
|  | pH 5 | 6,08 | 3,90 | 20,5 |
|  | pH 8 | 5,07 | 4,01 | 21,1 |

^a^Parameters from fitting the data in Figure S4 to the linear extrapolation method

^b^Calculated from $\Delta G_{U}=C_{M}\cdot\left\langle m_{app} \right\rangle$

**Figure S1 – Crystal structure of EXG:CBM^QQQW,Δ2-5^**. A) Crystal packing in the solved structure. B) Validation summary from wwPDB comparing the solved structure to other deposed structures. C) Alignment of the eight chains in the asymmetric unit.

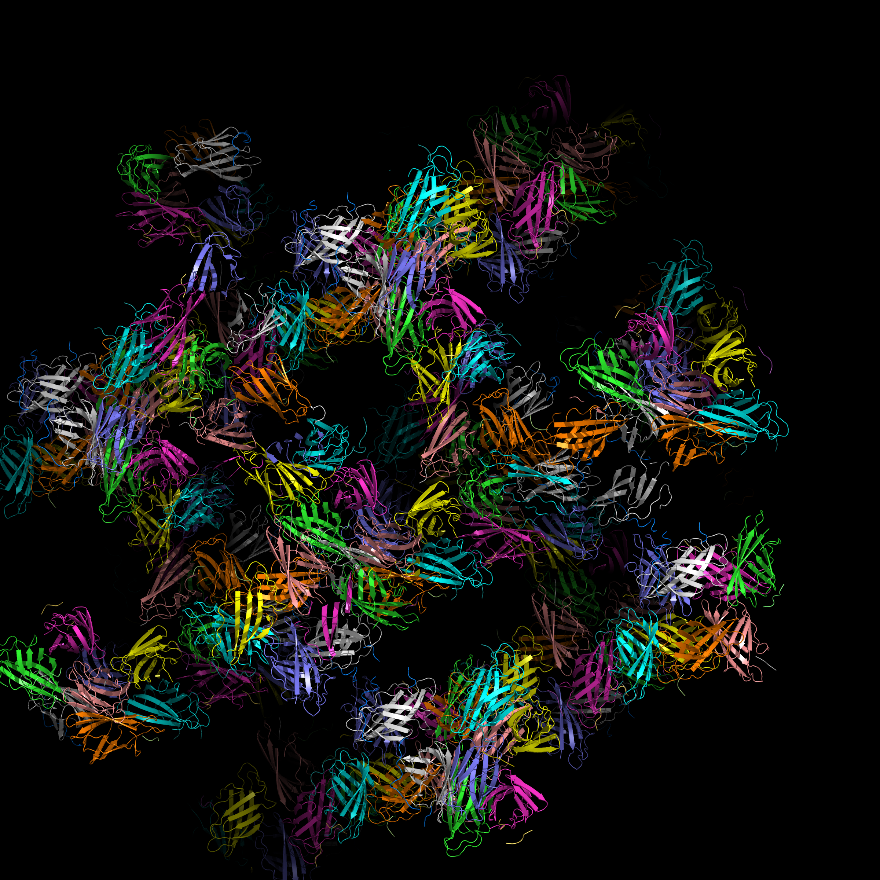

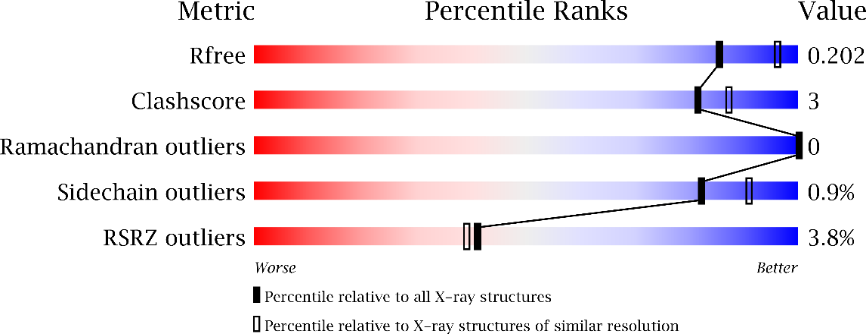

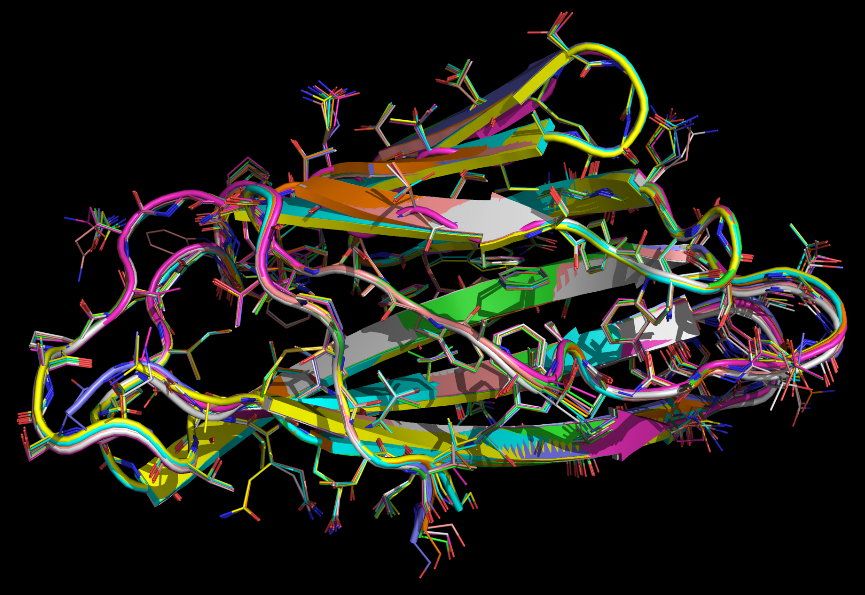

**A**

**B**

**C**

**Figure S2 - pH titration followed by NMR chemical shifts**. Selected chemical shifts as function of pH. Histidine and C-terminus titrations could be followed entirely by ^13^C-chemical shifts, while the ^13^C^γ^-shifts of Asp often disappeared at low pH. Therefore, Asp pK_a_-values were fitted globally from ^13^C^γ^ and amide signals. Color legend from Asp C^γ^ titration curve also applies to Asp H and Asp N nuclei.

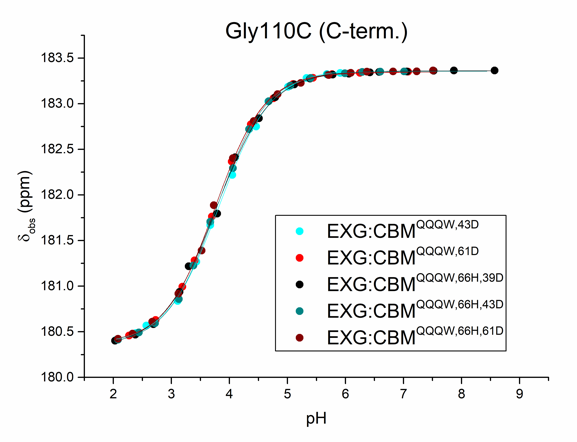

**Figure S3 – pK_a_ prediction from the protonation state sampling by CpHMD.** ***A)*** Titration curves for the three single aspartate EXG variants. The degree of proton dissociation at various pH was fitted to the Hill equations. ***B)*** Titration curve of (**top**) the aspartate residues (D39, D43, and D61) in the double titratable EXG variants fitted to the Henderson-Hasselbalch equation (solid line. Titration curve of (**bottom**) of the histidine residue (His66) in the double titratable EXG variants fitted to the Henderson-Hasselbalch equation (solid line). For Asp39 and His66 in the presence of Asp39 the data cannot be fitted to the Henderson-Hasselbalch equation to obtain pK_a_ values and Hill coefficients.

**A**

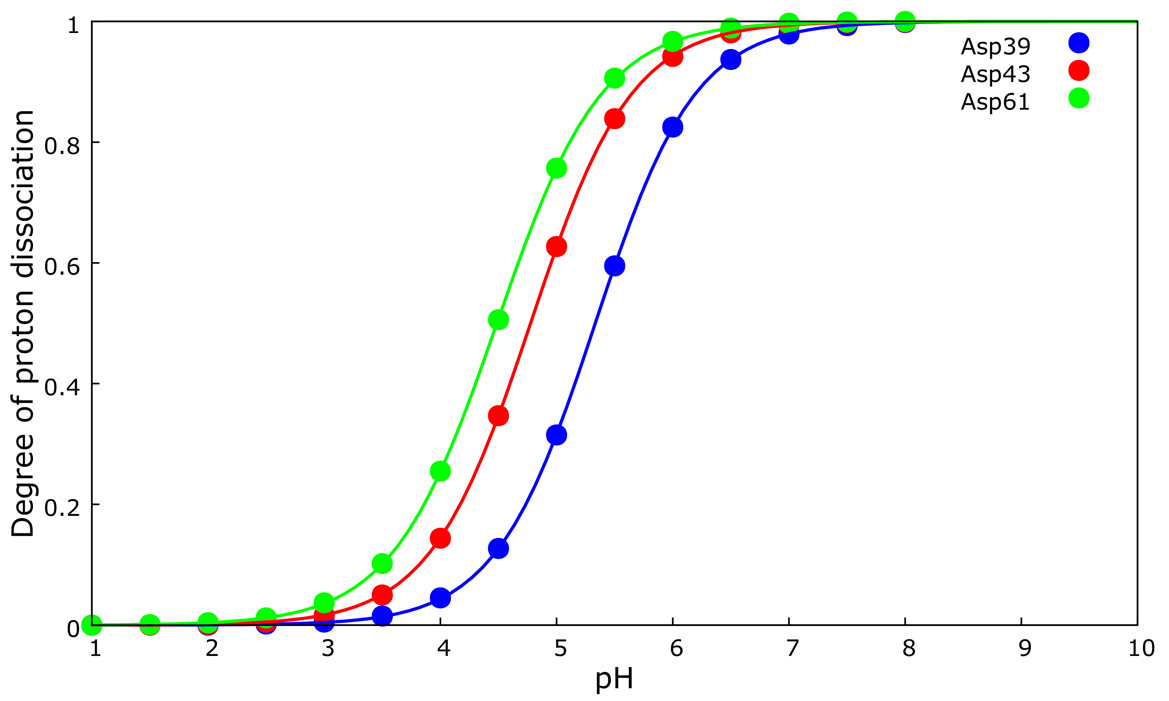

**B**

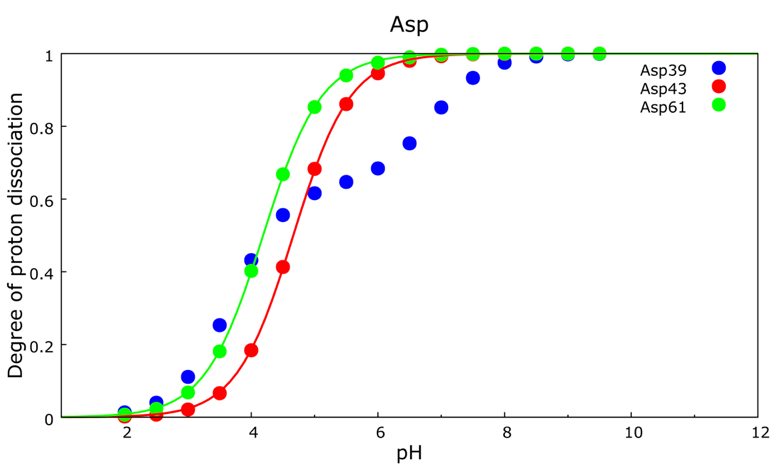

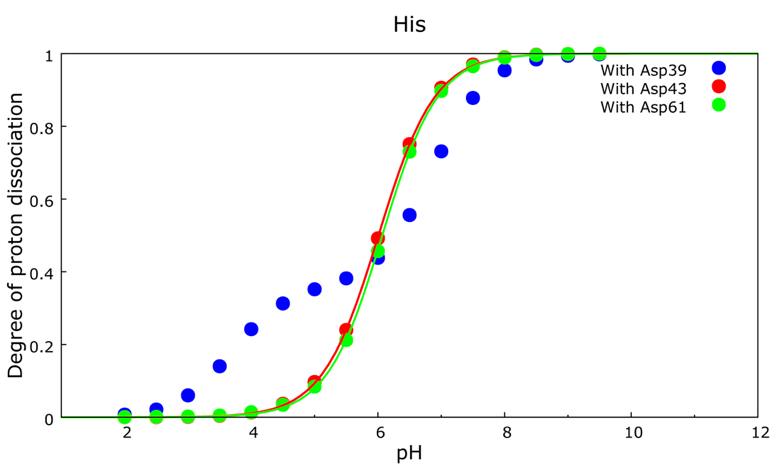

**Figure S4 – Urea induced unfolding data.** Raw data from fluorescence measurements. Lines represent global fits to the linear extrapolation method.

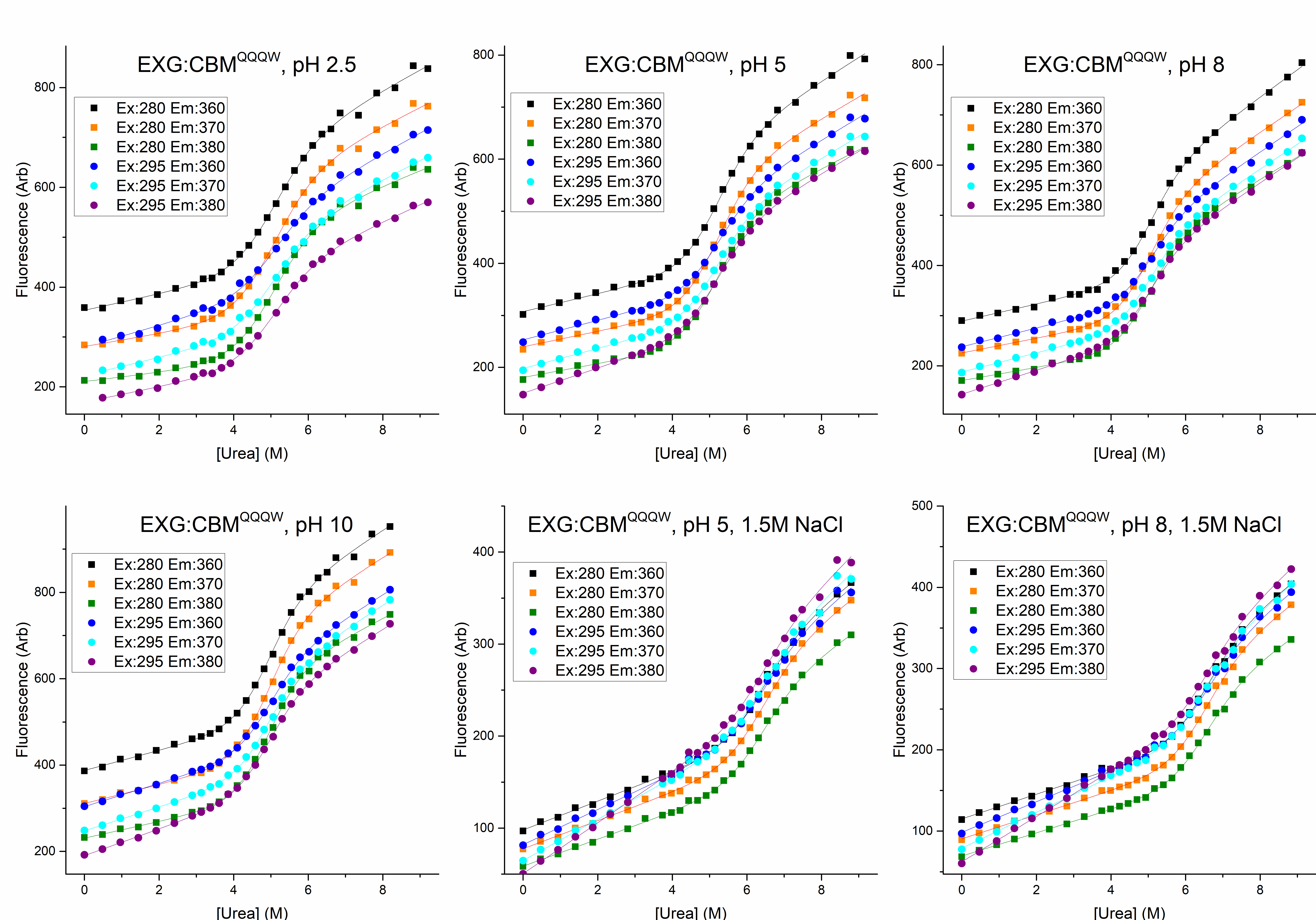

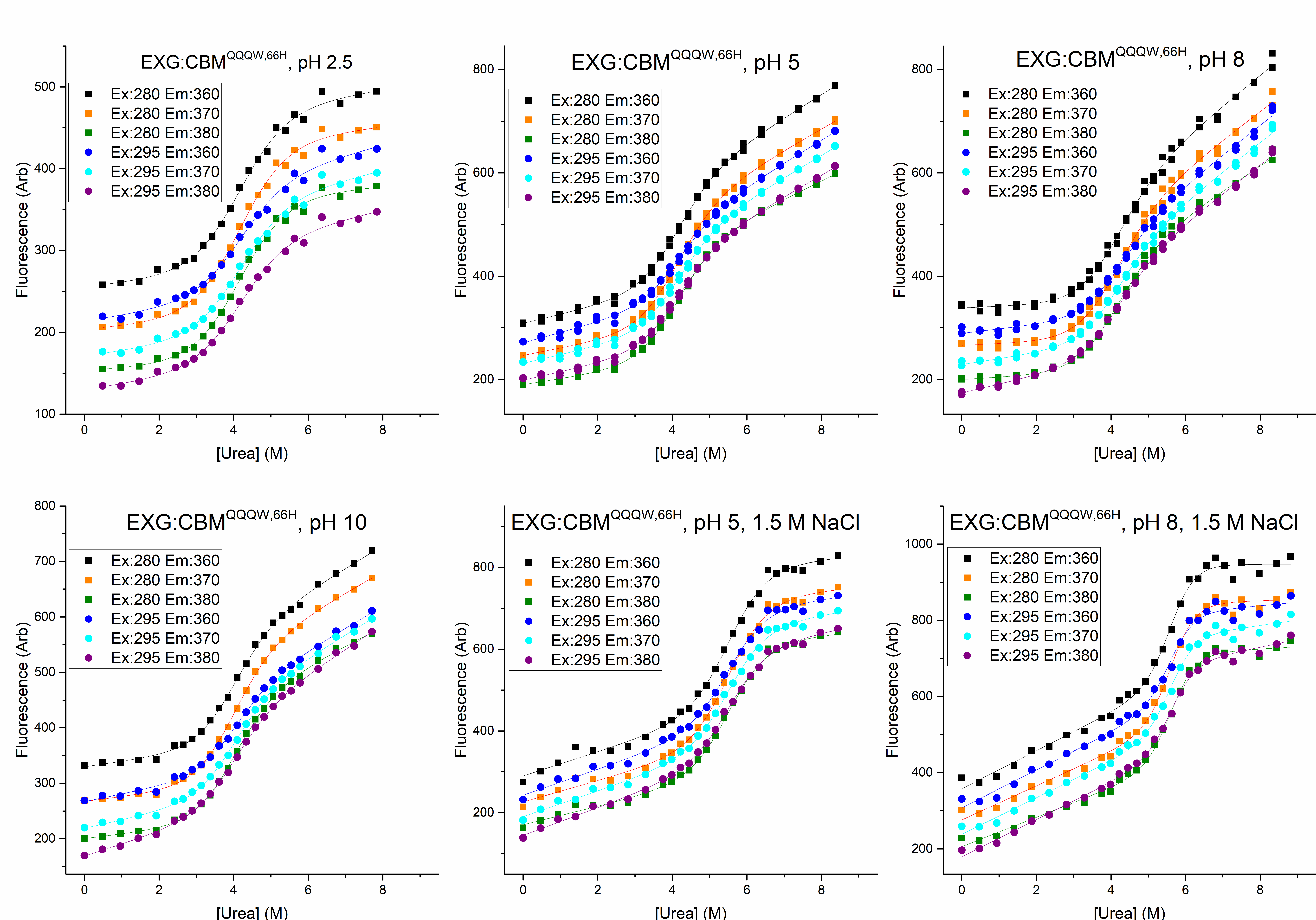

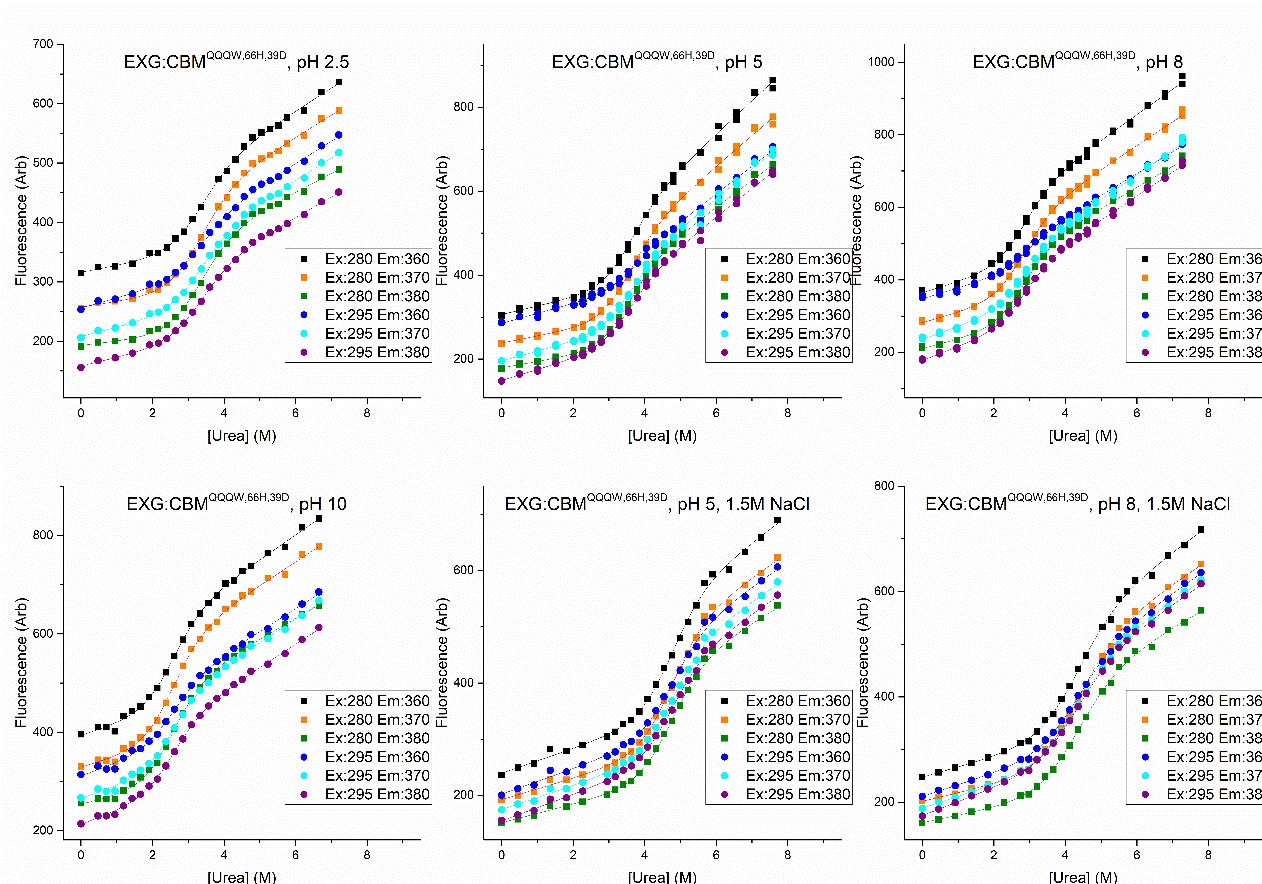

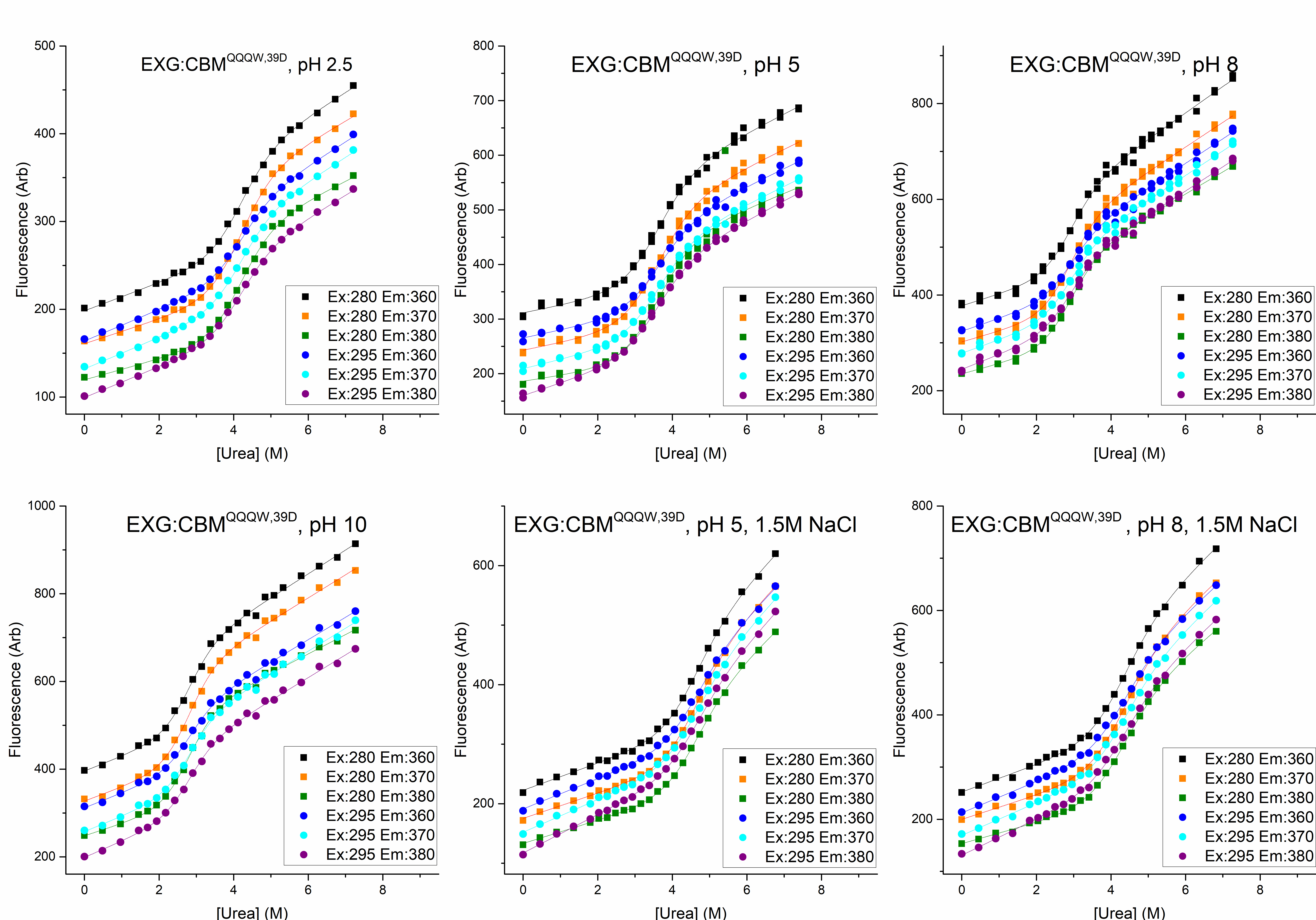

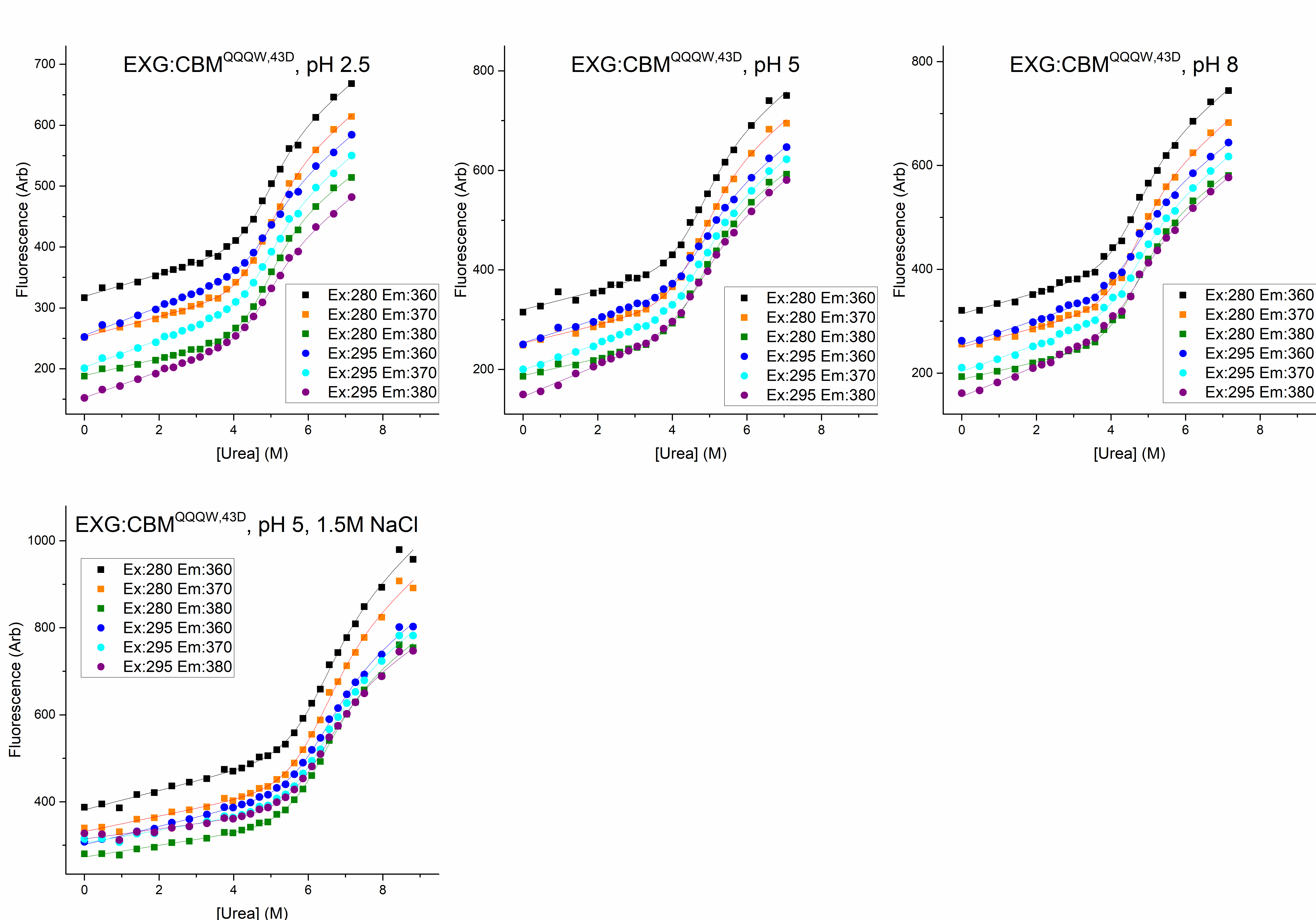

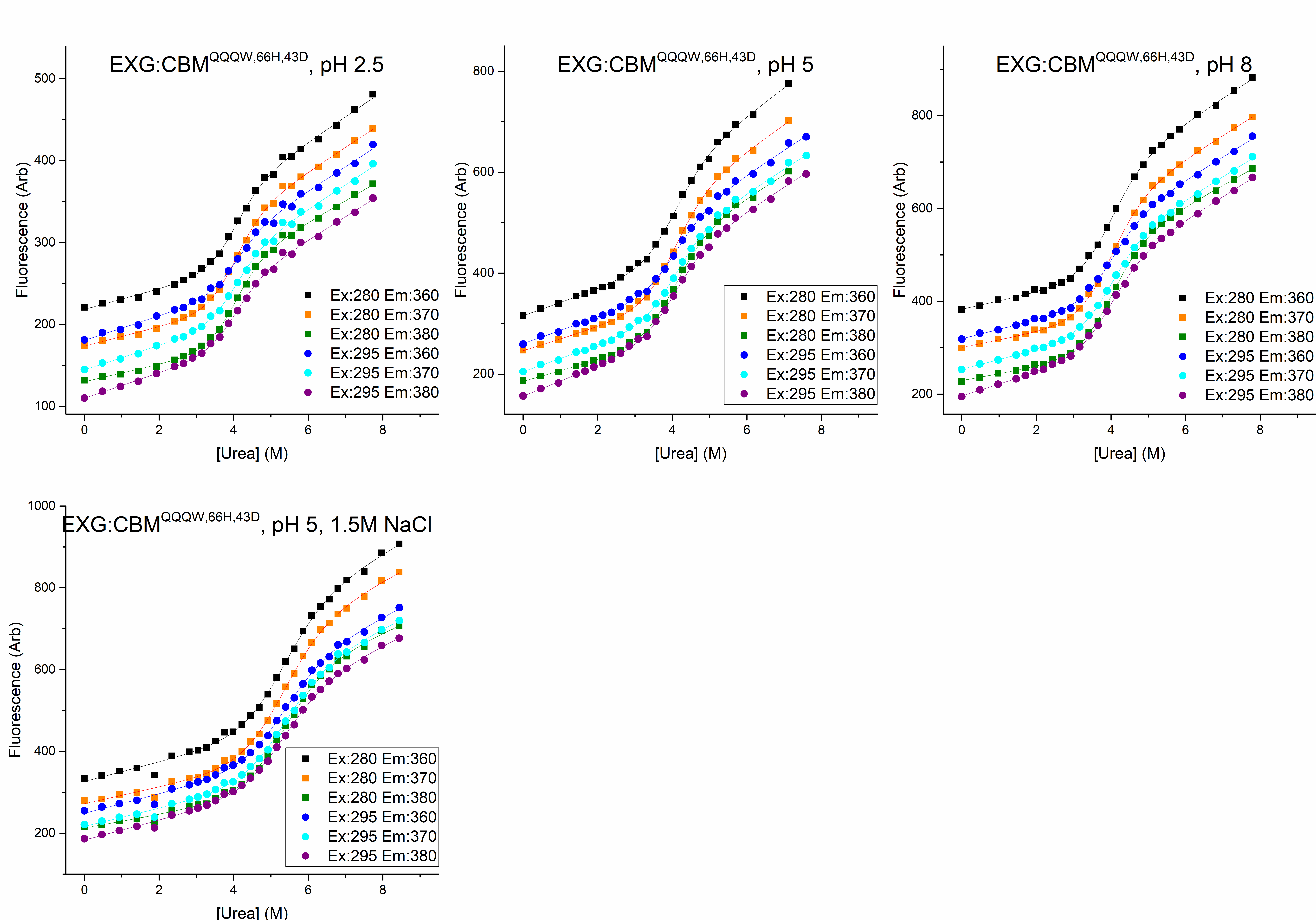

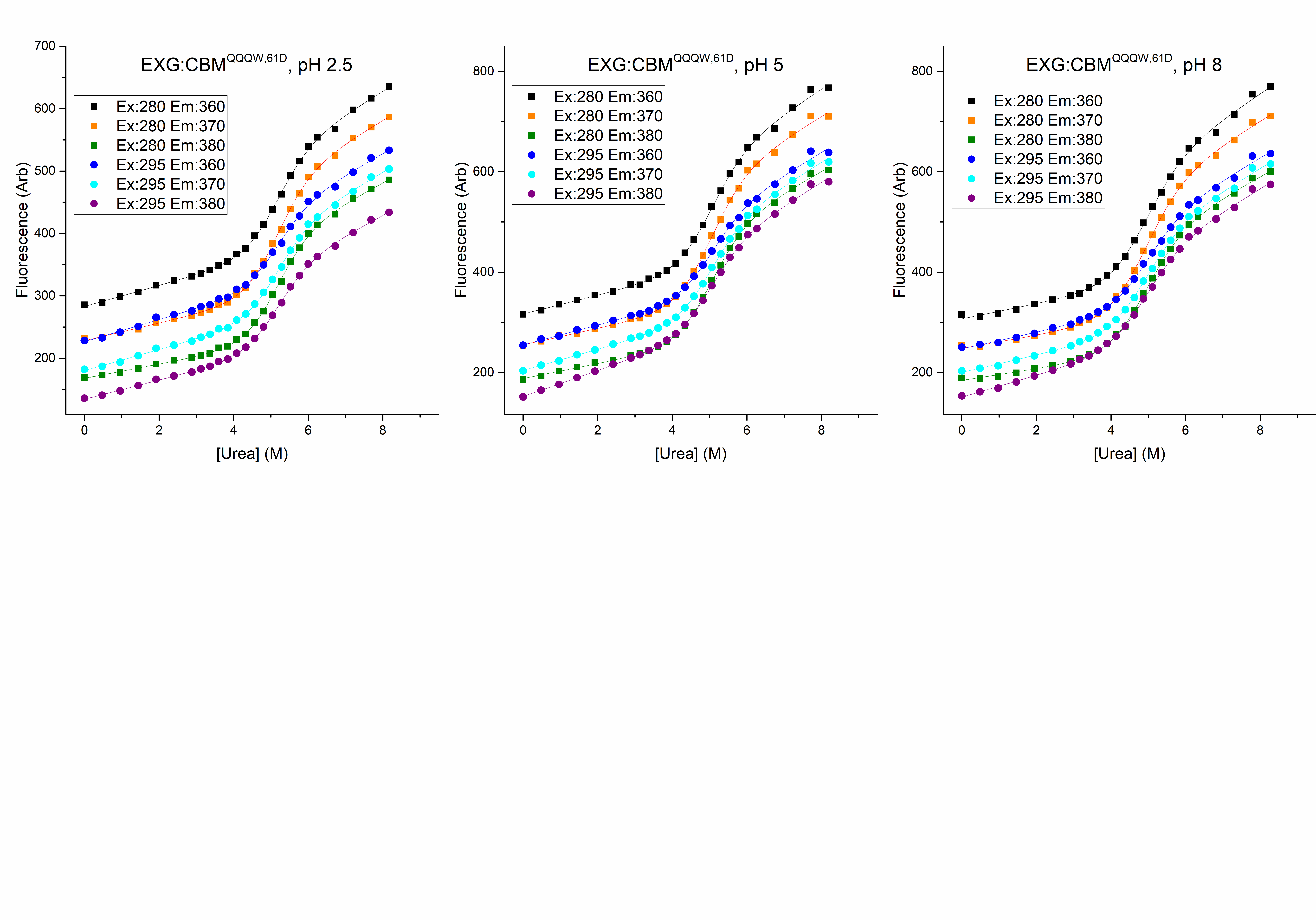

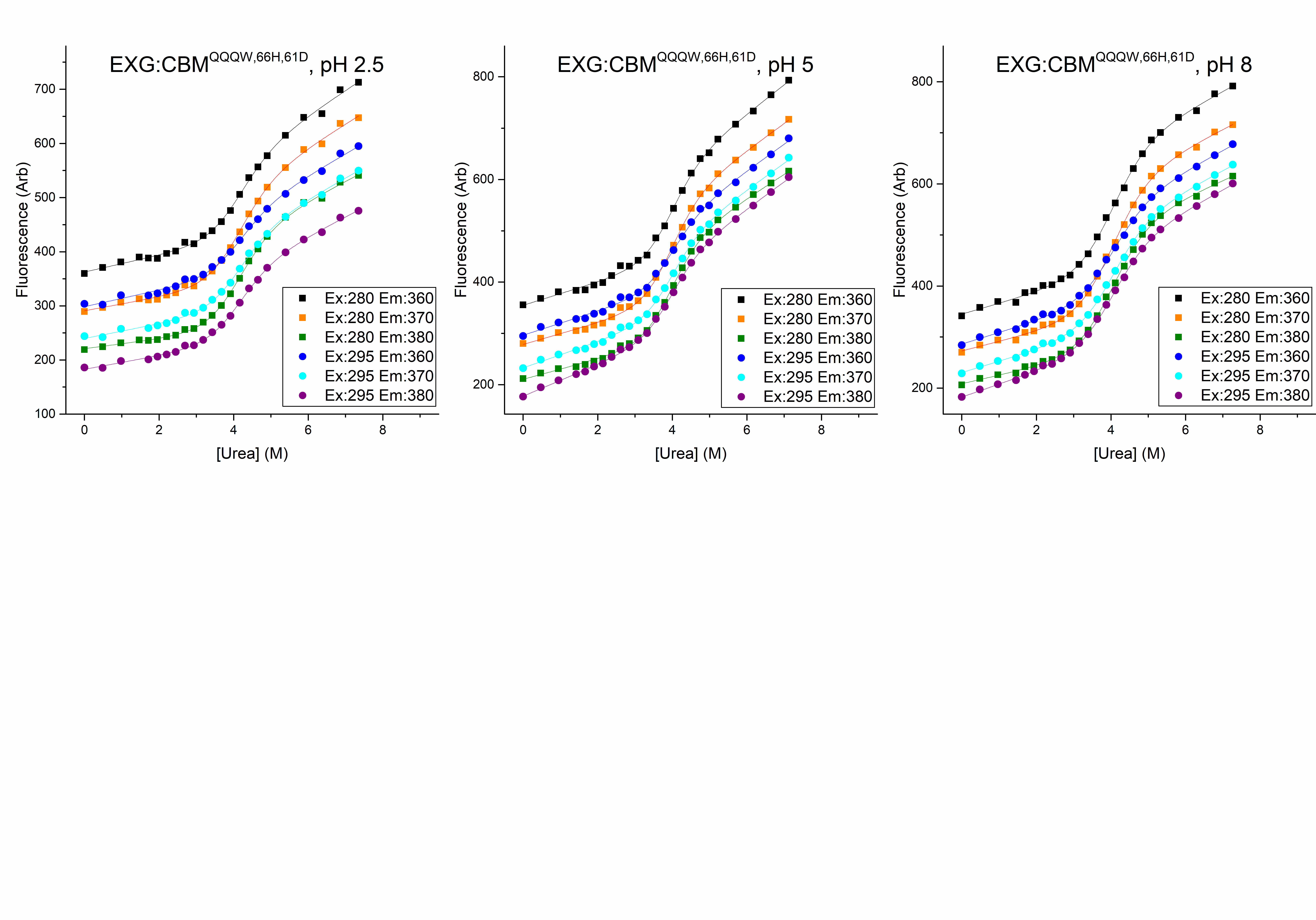

**Figure S5 – ^15^N-HSQC spectra of the investigated EXG:CBM variants.** The colors correspond to EXG:CBM^QQQW,43D^ (green), EXG:CBM^QQQW,61D^ (orange), EXG:CBM^QQQW,66H,39D^ (red), EXG:CBM^QQQW,66H,43D^ (blue), and EXG:CBM^QQQW,66H,61D^ (black). The spectra were measured at pH ~4.

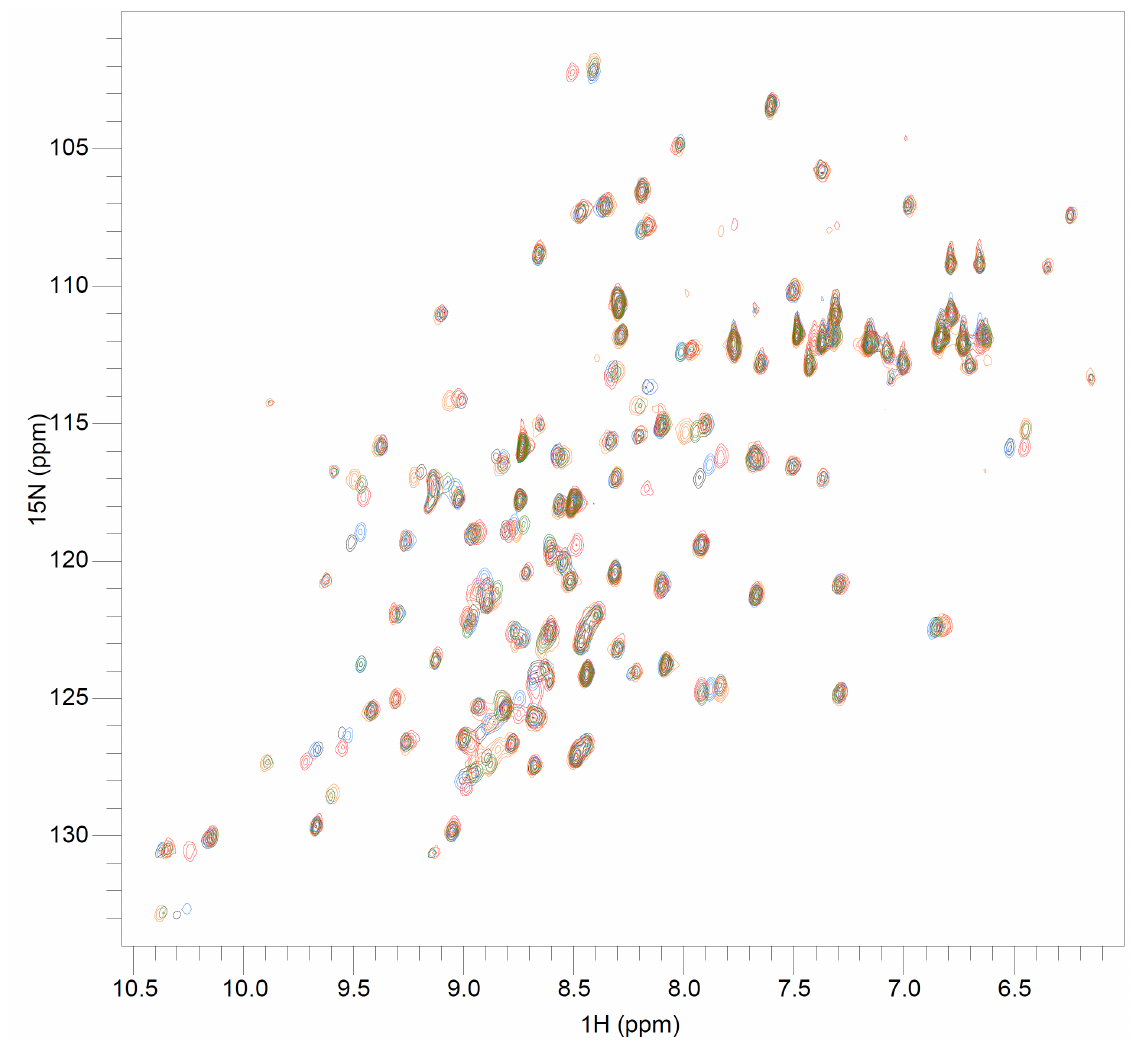

**Figure S6. Double-mutant cycles**.

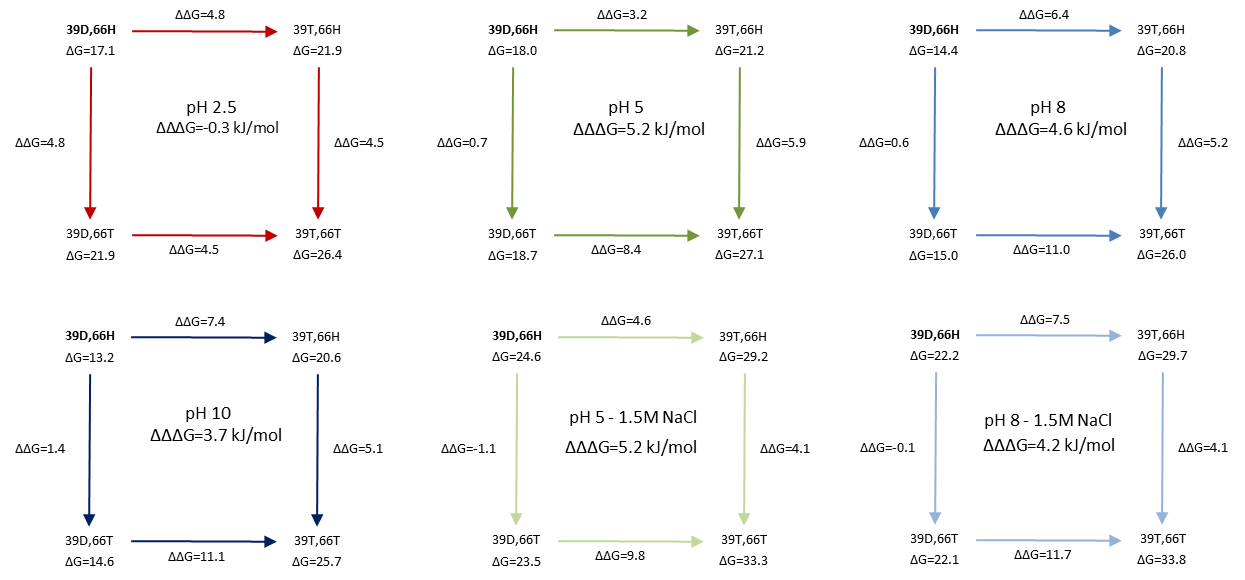

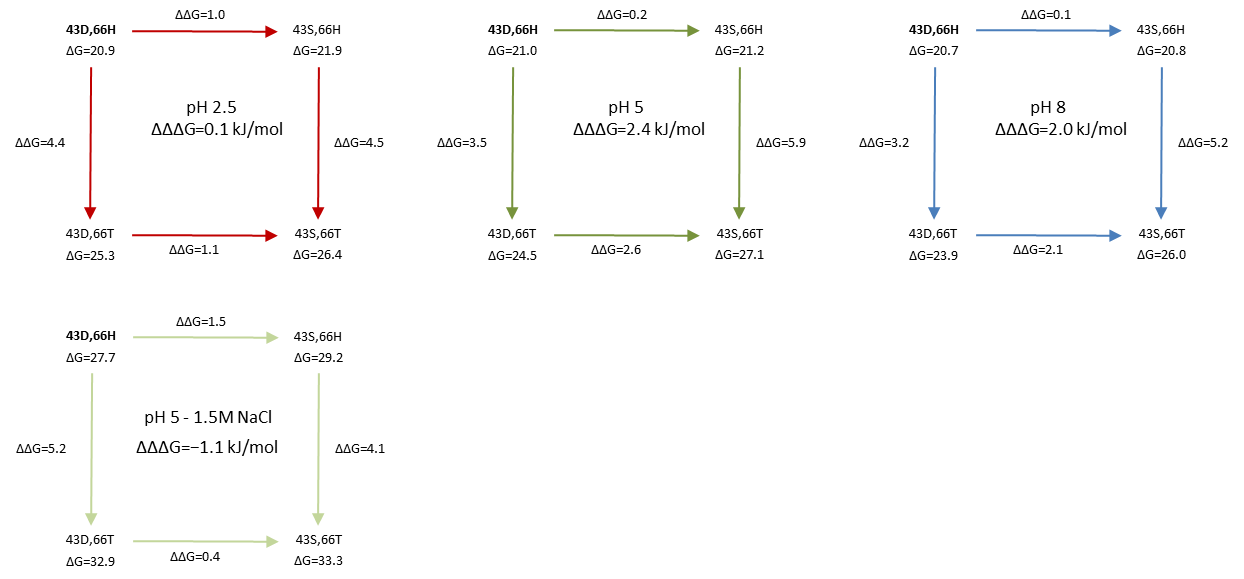

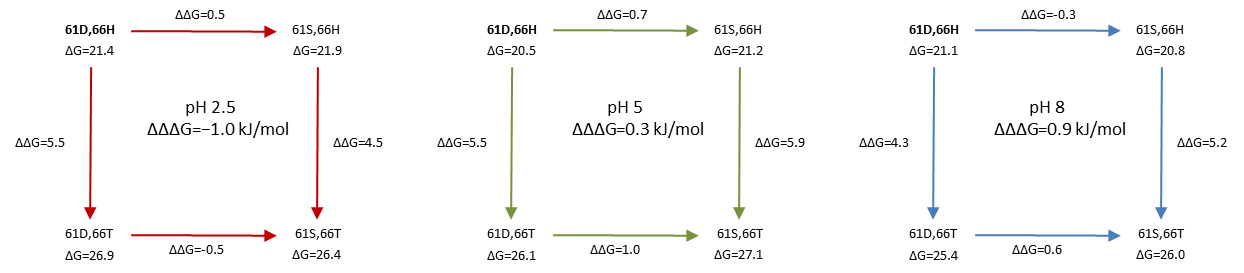

**Figure S7 – Rotamer population of Asp and His in double titratable EXG:CBM variants.** Histograms of the sampled χ_1_ dihedrals of Asp and His residues in the EXG:CBM^QQQW,39D,66H^ (top row), EXG:CBM^QQQW,43D,66H^ (middle row) EXG:CBM^QQQW,61D,66H^ (bottom row) from CpHMD at pH 2.0, 5.0 and 9.0.

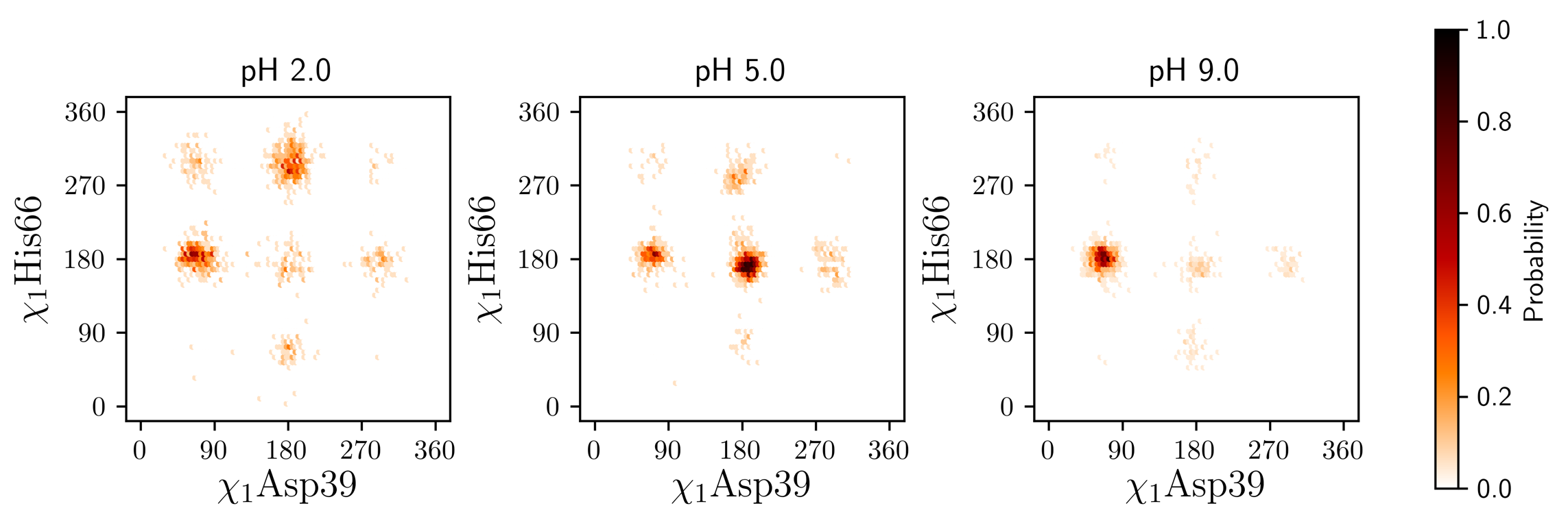

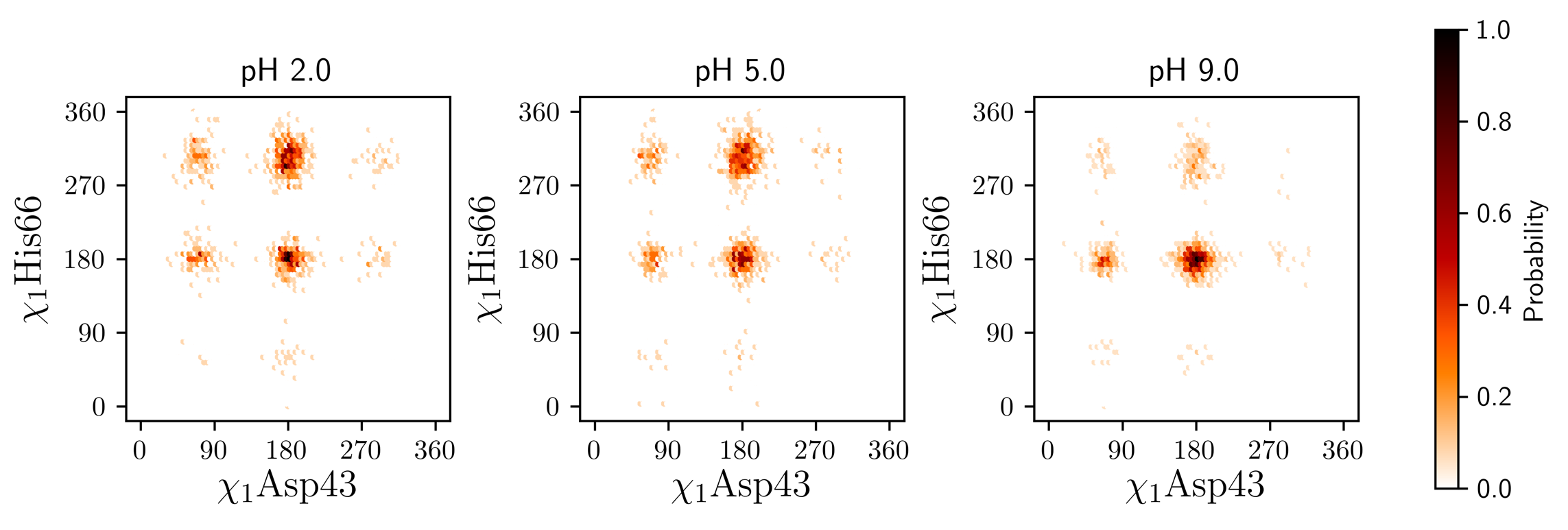

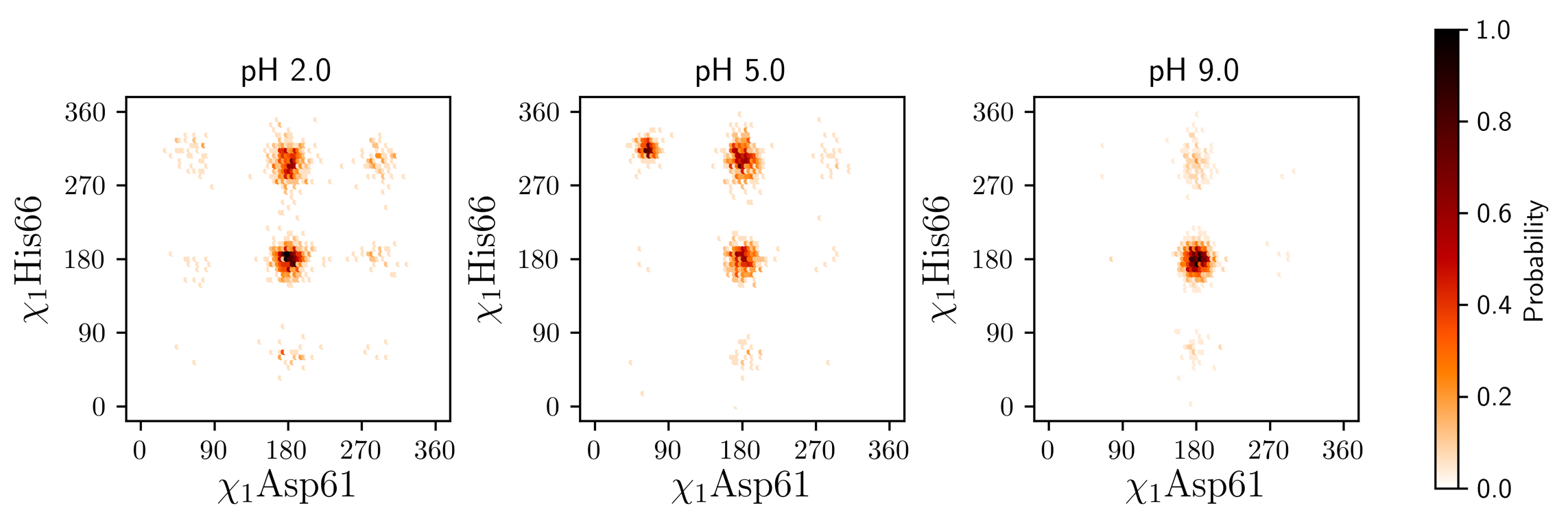

**Figure S8 – Rotamer population of Asp and His in double titratable EXG:CBM variants.** Histograms of the sampled χ_1_ dihedrals of Asp in EXG:CBM^QQQW,39D^ (left), EXG:CBM^QQQW,43D^ (middle) EXG:CBM^QQQW,61D^ (right) from CpHMD at pH 0.5 (blue) and 8.0 (orange).

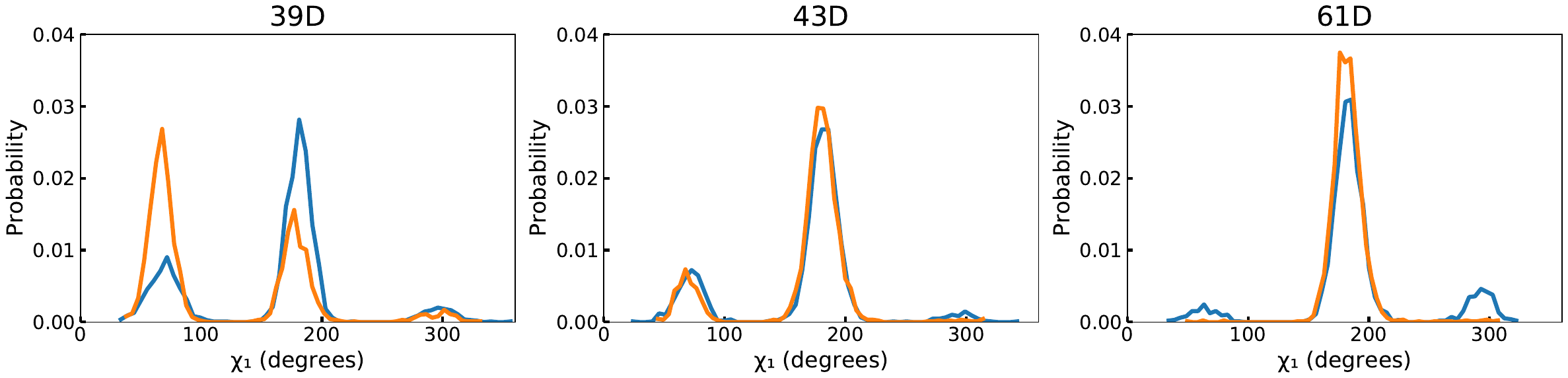

**Figure S9 – Thermodynamic cycle for estimating pK_a_ shifts from stability data.** Thermodynamic cycle relating the energy of deprotonation the residue *i (*$\Delta G_{deprot}^{i}$) in the folded (F) and unfolded (U) state of a protein, to the residues contribution to protein stability ($\Delta G_{U}^{i}$) at pH where the residue is fully protonated (p) and fully deprotonated (dp) respectively. Figure modified from (Bosshard et al, 2004)

$$\Delta G_{deprot}^{F,i}=2.3RT pK_{a}^{F,i}$$

F_dp_

**F**_p_

$$\Delta G_{U}^{dp,i}$$

$$\Delta G_{U}^{p,i}$$

U_p_

U_dp_

$$\Delta G_{deprot}^{U,i}=2.3RT pK_{a}^{U,i}$$

Deprotonation of folded protein

Deprotonation of unfolded protein

Unfolding of protonated protein

Unfolding of deprotonated protein

From Figure S7 it follows that:

Eq. S3

Eq. S2

Eq. S1

$$\Delta G_{deprot}^{F,i}+\Delta G_{U}^{dp,i}=\Delta G_{U}^{p,i}+\Delta G_{deprot}^{U,i}$$

$$\Delta G_{U}^{dp,i}-\Delta G_{U}^{p,i}=2.3RT \left( pK_{a}^{U,i}-pK_{a}^{F,i} \right)$$

$$\Delta pK_{a}^{U-F,i}=(\Delta G_{U}^{dp,i}-\Delta G_{U}^{p,i})/2.3RT$$

In a protein with multiple charged residues, the total change in stability, $\Delta\Delta G_{U}$, integrated over the entire pH range is thus related to difference in pK_a_-values between folded and unfolded state of all its residues (Bosshard et al, 2004):

Eq. S4

$$\Delta\Delta G_{U}=\Delta G_{U}^{dp}-\Delta G_{U}^{p}=2.3RT\sum_{i=1}^{n} (pK_{a}^{U,i}-pK_{a}^{F,i})$$

The total change in protein stability between two pH is determined by the sum of the contributions from all charged residues (Eq. S4). In EXG:CBM^QQQW,39D^, there are only three potential charges: Asp39 and the C- and N-terminus. The effects of the C- and N-termini on $\Delta\Delta G_{U}$ can be calculated from $\Delta G_{U}^{dp}-\Delta G_{U}^{p}$ in EXG:CBM^QQQW^. Subtracting the found energy difference from the $\Delta\Delta G_{U}$ of EXG:CBM^QQQW,39D^ leaves the contribution of Asp39, assuming the contribution of the C- and N-termini are the same in the two proteins. This energy was used in Eq. S3 to estimate the pK_a_ difference of Asp39 in the folded and unfolded state. Stability data from pH 2 (Asp39 fully protonated) and pH 8 (Asp39 fully deprotonated) were used for the calculation.
